# PPR9 mediates mitochondrial *nad* transcript maturation required for complex I biogenesis and early plant development in Arabidopsis

**DOI:** 10.64898/2025.12.04.692323

**Authors:** Ella Kobaivanov, Marketa Kitel, Roei Matan, Ron Mizrahi, Noa Carmi, Oren Ostersetzer-Biran

**Author notes:** These authors contributed equally to this work (Ella Kobaivanov, Marketa Kitel).

## Abstract

The biogenesis of plant mitochondria and their respiratory machinery relies on the transcription and proper processing of primary transcripts into functional, mature organellar RNAs. This includes the removal of introns that reside within many essential organellar genes, an essential step in mitochondria gene-expression that relies on the coordinated action of nuclear-encoded RNA-binding cofactors, such as members of the pentatricopeptide repeat (PPR) protein family. Here, we report the analysis of PPR9 (At1g03560), a mitochondrial P-type PPR protein originally identified in genetic screens for essential protein cofactors. Loss of PPR9 function results in embryonic arrest at early stages of seed development. Nevertheless, homozygous *ppr9* mutants can be maintained using a modified embryo-rescue method. The rescued *ppr9* mutants display delayed germination and growth retardation associated with mitochondrial dysfunctions. Molecular analyses of *ppr9* plantlets reveal that PPR9 is required for the splicing of several mitochondrial transcripts encoding respiratory subunits, including *nad2* intron 3 and *nad7* introns 1 and 2. Accordingly, defects in the processing of *nad2* and *nad7* pre-RNAs in *ppr9* mutant plantlets impair the biogenesis of respiratory complex I (CI), disrupt OXPHOS activity, and consequently affect plant’s growth and development. Together, these findings identify PPR9 as a key regulator of mt-RNA maturation in Arabidopsis, enabling CI biogenesis and cellular energy metabolism, and further highlight the roles of nuclear-encoded RNA-processing factors in coordinating mitochondria functions and (early) plant growth and development.

**Significance statement:** Our study identifies PPR9 as a key mitochondrial RNA-processing factor required for the splicing of multiple group II introns that reside within genes encoding subunits essential for respiratory complex I biogenesis in Arabidopsis. By linking nuclear control of RNA maturation to mitochondrial function and plant development, this work emphasizes a critical layer of coordination between organellar gene expression, energy metabolism, and developmental regulation in land plants.

## Introduction

Mitochondria are essential organelles that function as hubs for diverse metabolic activities, including as key sites of energy production, and in signaling pathways in eukaryotic cells [9, 62, 65]. Mitochondria in plants contribute to key developmental processes, such as developmental regulation and stress responses. As remnants of an ancestral α-proteobacterial endosymbiont, mitochondria retain a bacterial-related genome (mtDNA, mitogenome), with intrinsic transcriptional and translational machineries [27]. However, the vast majority of the organellar proteins, as well as some tRNAs, are encoded by nuclear loci and imported into the mitochondria (for recent reviews see e.g., Braun 2020, Møller et al. 2021). The assembly of the oxidative phosphorylation (OXPHOS) machinery requires a coordinated expression of both nuclear- and mitochondria-encoded proteins. These processes are essential for maintaining fictional respiration and cellular homeostasis, thereby enabling optimal plant growth and development.

Although all mitochondria are thought to share a common ancestor, they have diverged considerably across different eukaryotic lineages. Plant mitogenomes differ markedly from their animal counterparts in both size and structural complexity. Whereas animal mitogenomes are typically compact (∼17 kb) and conserved, those of land plants vary dramatically in size, ranging from tens of kilobases to over 10 Mb, and are further characterized by multipartite and highly recombinogenic structures [7, 27, 31, 33, 38, 47, 66, 90]. This genetic complexity is further evident in mitochondrial gene-expression, where plant organellar transcripts must undergo extensive post-transcriptional processing, including trimming, editing and splicing, to produce functional rRNAs, tRNAs, and mRNAs that can be translated into the correct organellar proteins [37, 92, 111].

A noticeable feature of plant mitogenomes is the high abundance of intron sequences that reside in many protein-coding genes, particularly within transcripts encoding subunits of the NADH dehydrogenase enzyme (i.e., complex I, CI), and also in genes encoding cytochrome c oxidase complex (CIV) subunits, cytochrome c maturation (CCM) factors, ribosomal protein biogenesis, but may reside in other genes as well [6, 11, 83, 111]. The majority of the organellar introns are identified as group II-type, an ancient group of catalytic ribozymes that are evolutionarily related to non-LTR retroelements, which also share common features with spliceosomal introns and telomerases [52, 109, 110]. Canonical group II RNAs are characterized by a conserved secondary structure comprising of six domains (D1-to-D6), which allow the folding of an active tertiary conformation that undergoes two sequential self-transesterification reactions required for splicing [59, 73, 109, 110]. Yet, for efficient excision *in vivo*, group II introns rely on their own intron-encoded maturase proteins (IEPs, or MATs). The splicing of model group II introns may also rely on additional factors, such as RNA helicases, which like the cognate MAT factors, likely assist in stabilizing the group II intron structure and thus promoting efficient RNA catalysis [32, 36, 52, 63, 64, 79, 83, 108]. Throughout evolution, the plant mitochondrial introns have diverged extensively in sequence and have often lost their intron-encoded proteins [64]. In several cases, the introns have fragmented into independently transcribed gene fragments that assemble *in trans* to facilitate exon joining [6, 64]. The sequence (and likely structural) alterations of the organellar introns were accompanied by the recruitment of nuclear-encoded cofactors, to facilitate the splicing of the non-canonical introns in plant mitochondria, and are further contributing to increased regulatory complexity of organellar gene expression. The plant organellar splicing factors belong to a diverse set of RNA-binding proteins, such as nuclear-encoded maturase-like proteins (nMATs), RNA helicases, CRM-related factors, mTERFs, PORR-domain proteins, and members of the greatly expanded pentatricopeptide repeat (PPR) protein family in plants, which are in the focus of this study [11, 37, 63, 64, 92, 111].

The PPR factors constitute one of the largest protein families in land plants [2, 14, 76, 81, 93, 94, 97, 102]. These are characterized by tandem repeats of a degenerate 35-amino acid motif (with a helix-turn-helix structure), enabling a modular and sequence-specific recognition of single-stranded RNA, where each PPR motif is postulated to bind a specific ribonucleotide (see e.g., [2, 30, 86]. Many of the plant PPR proteins contain putative mitochondria localization signals, where they are expected to regulate diverse aspects of organellar RNA metabolism, including RNA processing and trimming, transcript stabilization, RNA editing (primarily C-to-U deamination), and the splicing of transcripts containing group II introns [34, 37, 54, 81, 92, 111]. In addition, accumulating data from recent studies suggest that some PPR proteins may also participate in the regulation of mitochondrial translation [26, 77, 104].

Despite substantial advances in understanding the RNA-recognition code and target specificity [2, 15, 35, 60, 75], as well as the specific roles of PPR proteins in RNA editing [94], the molecular mechanisms by which PPR proteins facilitate splicing remain poorly understood. Here, we report the functional characterization of PPR9 (At1g03560), a P-class PPR protein that is essential for early embryogenesis in *Arabidopsis thaliana*. Experimental analyses demonstrate that PPR9 is required for the excision of several mitochondrial group II intron (i.e., *nad2* intron 3 and *nad7* introns 1 and 2). Impaired splicing of these introns in *ppr9* mutant-plants impairs holo-CI biogenesis, leading to severe physiological defects. The effects of PPR9 disruption on organellar biogenesis and the physiology of embryo-rescued Arabidopsis mutant-lines are described in detail.

## Results

### The *PPR9* (At1g03560) gene-locus encodes a P-type PPR protein predicted to localize to mitochondria

During the evolution of land plants, the mitogenomes (mtDNAs) have acquired remarkable structural plasticity, with extensive variation in genome size, architecture and expression patterns (especially at the RNA level) among species [4, 12, 27, 48]. To gain insights into the mechanisms of mt-RNA processing steps in plants, we screened a collection of publicly available *A. thaliana* T-DNA insertional mutants (i.e., The Arabidopsis Information Resource, TAIR, database) that reside in genes that encode putative mitochondria-targeted P-type PPR proteins. Among these, we noticed a heterozygous plant-line carrying a T-DNA insertion (SALK_004994) in the *AT1G03560* gene-locus (UniProt Q9LR67, or PPR9_ARATH), which failed to yield viable homozygous offspring. The deduced product of *PPR9_ARATH* gene (denoted here as PPR9) harbors 660 amino acids with a predicted topology of NH₂-151-(P)-P-P-P-P-P-P-P-P-P-P-P-P-P-19-COOH (Fig. 1a, Supplementary Fig. S1a), where ‘P’ denotes canonical PPR motifs [15, 107], while numbers correspond to amino acids regions that are not assigned to any defined domains. The first PPR shown in quotes represents a degenerated PPR motif, as predicted by the PPR code (https://ppr.plantenergy.uwa.edu.au) [15] and the AlphaFold servers. TargetP [20], UniProt [106] and SUBA4 [39] predict the presence of a 53-amino acid mitochondrial targeting peptide region (mtTP) at the N-terminus of PPR9 (Fig. 1a). *In silico* 3D modeling of Arabidopsis PPR9 with AlphaFold [42] further indicate a canonical superhelical solenoid fold (Fig. S1), containing an internal basic groove within the molecule that is consistent with RNA-binding regions described for PPR proteins [29, 30, 87].

**Figure 1.**
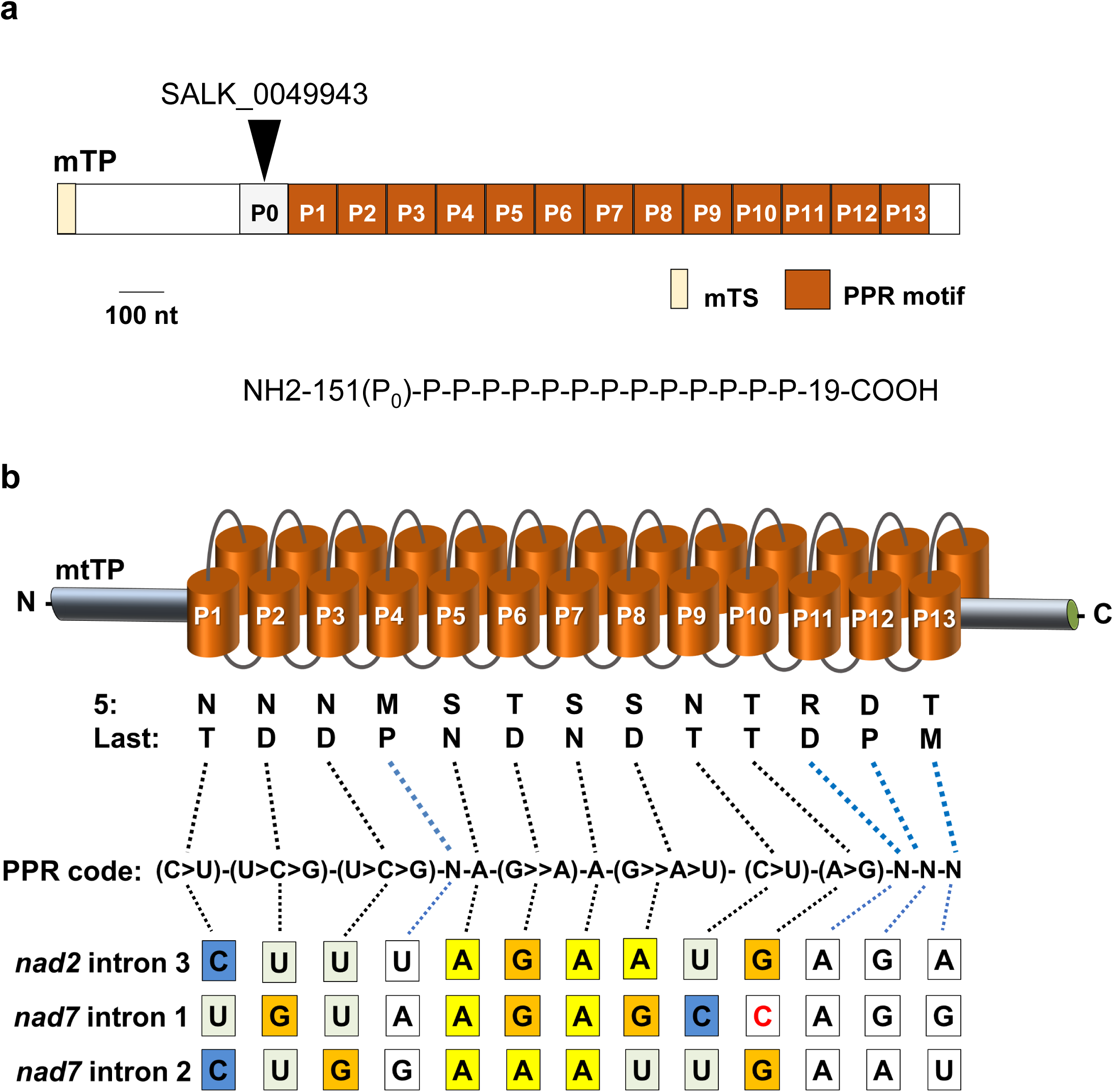
The *AT1G03560* gene-locus (*PPR9*) encodes a putative mitochondria-localized PPR protein. (a) Schematic representation of the *PPR9* (*AT1G03560*, UniProt Q9LR67) gene structure. *In silico* analysis suggests the presence of 13 conserved P-class PPR motifs (P1-P13), as well as an additional putative PPR motif (P?) located near the N-terminus of the protein. Numbers correspond to amino acid residues not assigned to any defined motif. The predicted mitochondrial targeting signal (mTS) and the T-DNA insertion site in the SALK_004994 line (*ppr9-1*) within the PPR9 coding region is indicated above the gene structure figure. (b) Prediction of putative RNA-binding sites of PPR9. The fifth and last amino acids of each PPR repeat are indicated below the corresponding motifs. The predicted RNA-recognition code is shown beneath the figure. ‘N’ denotes PPR motifs with unassigned nucleotide specificity. Postulated RNA-binding sites within *nad2* intron 3 and *nad7* introns 1 and 2 (Table S1) are illustrated, with boxes color-coded as follows: U (light green), C (blue), G (orange), A (yellow), and unassigned bases (white). Red color indicates mismatches between the predicted PPR code and target nucleotides.

PPR proteins are characteristically defined act as sequence-specific RNA-binding factors [2, 30, 86, 92, 94], in which the RNA recognition is based on a combinatorial code generally defined by amino acids found at positions 5 and 35 (or the last residue in non-canonical domains) of each PPR motif that together define the nucleotides binding site [1, 15, 99]. Based on the RNA-binding code, several PPR proteins exhibit predicted binding site that accurately reflect the specific RNA target(s) validated through genetic analyses. In other cases, however, the predicted RNA-binding sites can be less defined or insufficient to accurately indicate the native target region. Likewise, when the PPR code was applied to the 13 canonical motifs of PPR9 (Figure 1b), it yielded a very broad range of potential binding sites within the mtDNA of *Arabidopsis thaliana* [91]. In total, 203 putative PPR9 recognition sites (excluding non-coding/intergenic regions) were indicated in Arabidopsis mitogenome [5, 91]. These were distributed across multiple gene regions: 19 sites are located within 5′ and 3′ untranslated (UTR) regions, 51 sites within exonic sequences, and 54 sites are found within intron sequences (Table S1). This ambiguity emphasizes the need for genetic and biochemical assays to elucidate the molecular function(s) of PPR9, by establishing its intracellular location(s), the RNA targets and roles in gene expression, *in vivo*.

### *PPR9* encodes a lowly-expressed mitochondria-localized PPR protein

Expression analysis using publicly available microarray and transcriptomic datasets [40, 57] indicates that PPR9 displays differential expression across developmental stages, with elevated transcript levels in embryonic tissues, young developing leaves, the shoot apex, apical root regions, and flowers (Fig. S2). Here, we further analyzed the intracellular location of PPR9 protein, *in vivo*. For this purpose, we created a fusion protein containing the first 82 amino acids of PPR9 in-frame with eGFP (N_82_-PPR9:GFP), and expressed it in protoplasts obtained from *P. patens* [100, 101, 105]. The confocal microscopy analyses showed that the PPR9-GFP signals appear as globular-shaped particles that co-localized with the MitoTracker marker, a mitochondrion specific fluorescent probe (Fig. 2). The confocal microscopy did not reveal any additional intracellular localization, such as chloroplasts (as indicated by chlorophyll autofluorescence), the nucleus, or the cytoplasm (Fig. 2, merged image). These results strongly support the *in silico* predictions as well as MS/MS data [25] that showed that PPR9 is targeted to the mitochondria, where it is postulated to function in post-transcriptional regulation of organellar gene expression and/or RNA metabolism [37, 92, 111].

**Figure 2.**
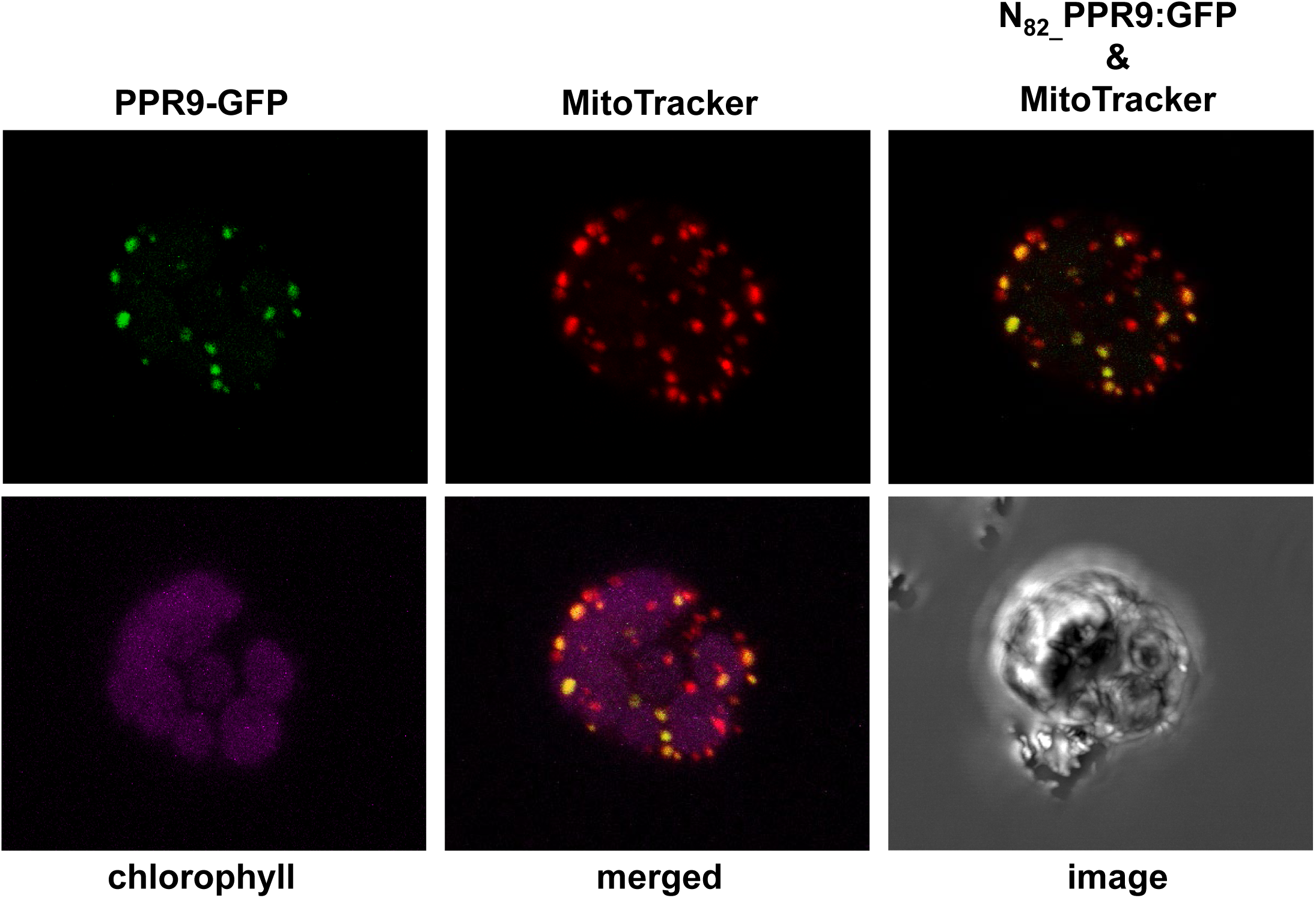
GFP localization analyses. Protoplasts were prepared from *P. patens* plantlets grown on sterile MS medium and transformed with the pCAMBIA-1302 vector containing a GFP sequence fused in-frame to the N-terminal region (246 bp) of the PPR9 (AT1G03560) coding sequence, under the control of CaMV 35S promoter and a NOS terminator sequence at the 3′ end. The figure shows confocal PPR9-eGFP fluorescence (green, top left panel) and MitoTracker (red, top middle panel) signals, a GFP-MitoTracker overlay (top right panel), chlorophyll autofluorescence (purple, bottom left panel), merged fluorescence channels (bottom middle panel), and a bright-field images (grayscale, bottom right panel).

### PPR9 functions are essential during early embryo development in *Arabidopsis thaliana*

Analysis of the SIGnAL database identified a single T-DNA insertion line in the *PPR9* coding region (SALK_004994), with the T-DNA located 536 nucleotides downstream of the AUG translation initiation codon (Fig. 1a). Genetic screening revealed that under optimal growth conditions (see Materials and methods), no homozygous plants could be obtained from the progeny of the heterozygous *ppr9* line (Fig. 3a). The heterozygous *ppr9* plants exhibited no visible abnormalities, suggesting that the complete loss of PPR9 functions result in embryonic developmental arrest or lethality. Siliques obtained from heterozygous *ppr9* plants (10 days after pollination) contained approximately one-quarter of white seeds (Fig. 3b), which later degenerated into shrunken and darkened seeds (Fig. 3a). Microscopic (Nomarski, DIC) analysis further revealed that green seeds contained fully developed embryos, whereas white seeds of the same progeny harbored embryos arrested at the torpedo (to walking-stick) stages (Fig. 3b). These phenotypes coincide with various other Arabidopsis mutants affected in mitochondria biogenesis and function [17, 69].

**Figure 3.**
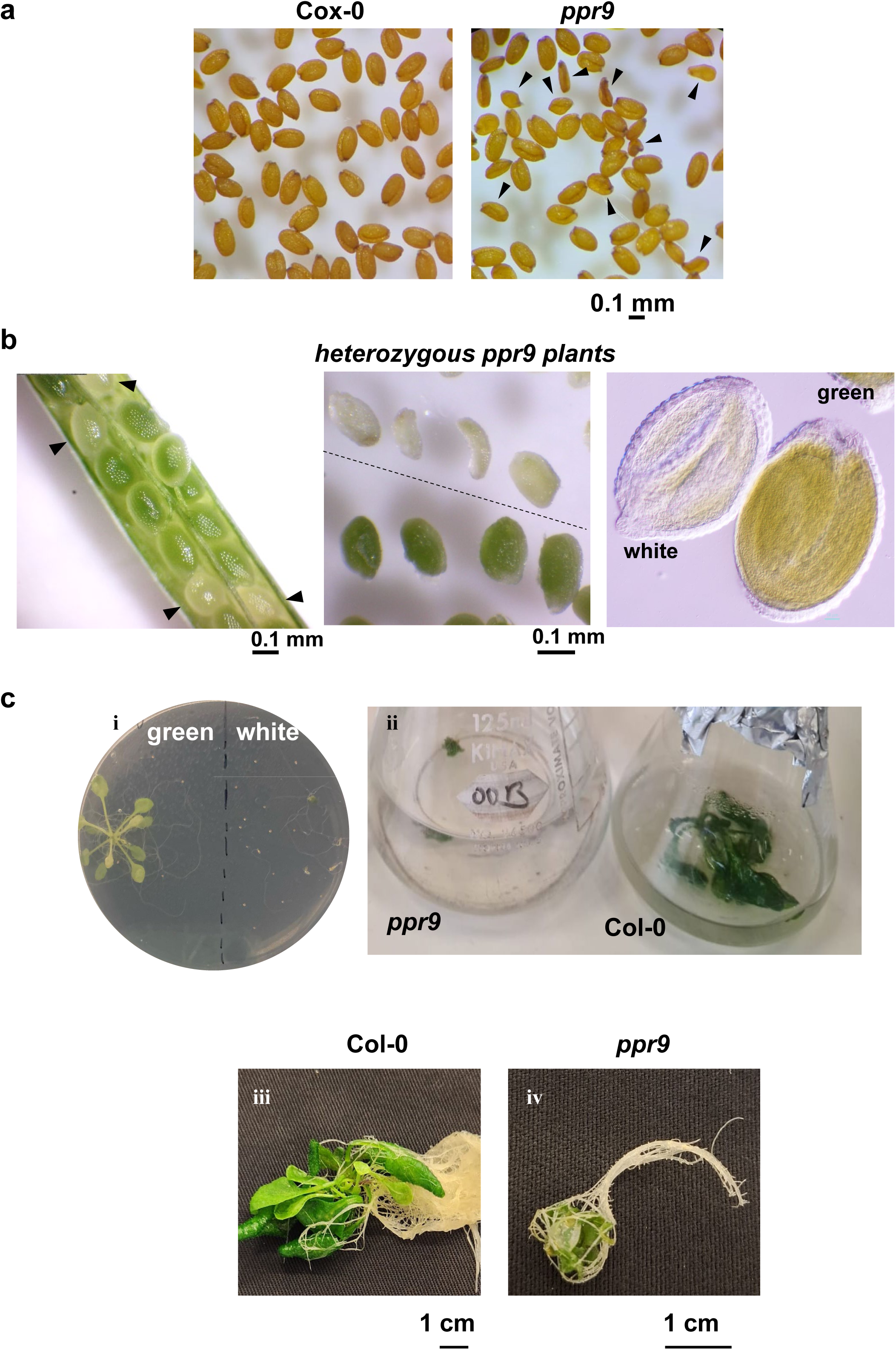
*ppr9* mutant phenotypes. Mature siliques obtained from heterozygous *ppr9* (SALK-004994) mutant line. (a) Seeds collected from mature siliques of wild type (Col-0) and heterozygous *ppr9* plants. Arrows indicate abnormal homozygous *ppr9* mutant seeds, which appear shrunken and darker. (b) Representative images of green (heterozygous or wild type embryos) and white (homozygous *ppr9* embryos) seeds observed in immature siliques of heterozygous *ppr9* plants (left panel). The morphology of embryos in green or white seeds (middle panel) were analyzed by differential interference contrast (Nomarski, DIC) microscopy (right panel). (c) Embryo-rescue procedure. Seeds collected under sterile conditions from surface-sterilized immature siliques of Col-0 or heterozygous *ppr9* plants were sown on MS-agar plates supplemented with 3% sucrose and vitamins (*i*), and grown under standard growth-chamber conditions. 2-month-old *ppr9* and Col-0 plantlets (growth stage L6) transferred to liquid MS medium supplemented with 3% sucrose and vitamins (*ii*). Homozygous *ppr9* plants exhibit abnormal morphology and developmental phenotypes, often displaying a bushy-like appearance (*iii*, *iv*), similar to that reported for other mutants affected in mt-RNA metabolism [3, 19, 88].

Embryo rescue techniques have been successfully applied to recover several Arabidopsis mutants exhibiting germination-defective phenotypes [23]. Equivalent approaches have been also used to rescue several mutants impaired in mitochondrial biogenesis or function in Arabidopsis, including *cod1* [19], *ndufv1* [49] *cal1/cal2* [18, 24], *nmat3* [88] and *msp1* [3]. Following our own laboratory strategy [3, 88], white seeds collected from green-mature siliques of heterozygous *ppr9* plants (10-12 days post-anthesis, DPA) were plated on MS-agar plates supplemented with 1% sucrose and a vitamin mix, and transferred to a controlled growth chamber (Fig. 3c). Under the *in vitro* conditions, between 20 to 30% of the white seeds germinated after about 1∼3 months (Fig. 3c-i). To increase biomass yield for downstream analyses (e.g., RT-qPCR, BN-PAGE), seedlings at the L6 stage [8] were transferred to liquid MS medium and cultured under the growth conditions specified in the ‘Plant material and growth conditions’ section, with gentle agitation (50-100 rpm). Most homozygous plantlets died within two to four weeks after germination or following transfer to liquid MS medium (Fig. 3c-i, Fig. S3). However, a small fraction (approximately 20% of those that germinated) survived and could be maintained for subsequent RNA, protein, and physiological analyses (Fig. 3c-ii, Fig. S3). PCR-based genotyping confirmed that while the green seeds from heterozygous *ppr9* plants were either wild type or heterozygous for SALK_004994 (Figs. 3c-ii, 3c-iii), while plantlets regenerated from white seeds (Figs. 3c-ii, 3c-iv) were homozygous for the *PPR9* mutant allele. As previously noted for various embryo-defective (*emb*) mutants [8], including those with defects in mitochondrial biogenesis (see e.g., [3, 17, 19, 49, 88], a growth retardation and phenotypic variability was also observed among the rescued *ppr9* plantlets (Figs. 3c-ii, 3c-iv, Fig. S3). No homozygous *ppr9* plantlets could be successfully established in soil, and none of the rescued plants analyzed in this study were able to complete a full life cycle or produce viable seeds.

### PPR9 functions in the processing of *nad2* and *nad7* RNA transcripts in *A. thaliana* plants

The accumulation (i.e., steady-state levels) of mitochondrial protein-coding transcripts in wild type and homozygous *ppr9* mutant lines was analyzed by RT-qPCR. Notably, the RNA profiles of the *ppr9* mutant plants (rescued white seeds) were compared with those of germinated wild type plantlets derived from the same immature embryos (green seeds) and propagated *in vitro* under the same growth conditions. These analyses indicated a ∼250-fold reduction in processed organellar transcripts corresponding to *nad2* exons ‘3’ and ‘4’ (*nad2* ex3-4) in *ppr9* mutant plants (Fig. 4a), suggesting impaired splicing of *nad2* intron 1. We further noticed reduced levels in transcripts corresponding to processed *nad7* exons ‘1’ and ‘2’ (*nad7* ex1-2; about 8-fold lower) and *nad7* exons ‘2’ and ‘3’ (*nad7* ex2-3; about 20-fold lower) in *ppr9*, suggesting to altered splicing in *nad7* introns 1 and 2, respectively, in *ppr9* mutant plants. The steady-state levels of various organellar transcripts in functionally complemented (embryo germinated) *ppr9/PPR9* plantlets, including *nad2* and *nad7* mRNAs, were comparable (i.e., differences were not statistically significant) to those in the wild type plants (Fig. 4b). Taken together, these data indicate that the loss of PPR9 function is associated with defects in the processing of *nad2* and *nad7* transcripts, and that the observed mt-RNA profiles are not attributable to the embryo-rescue procedure.

**Figure 4.**
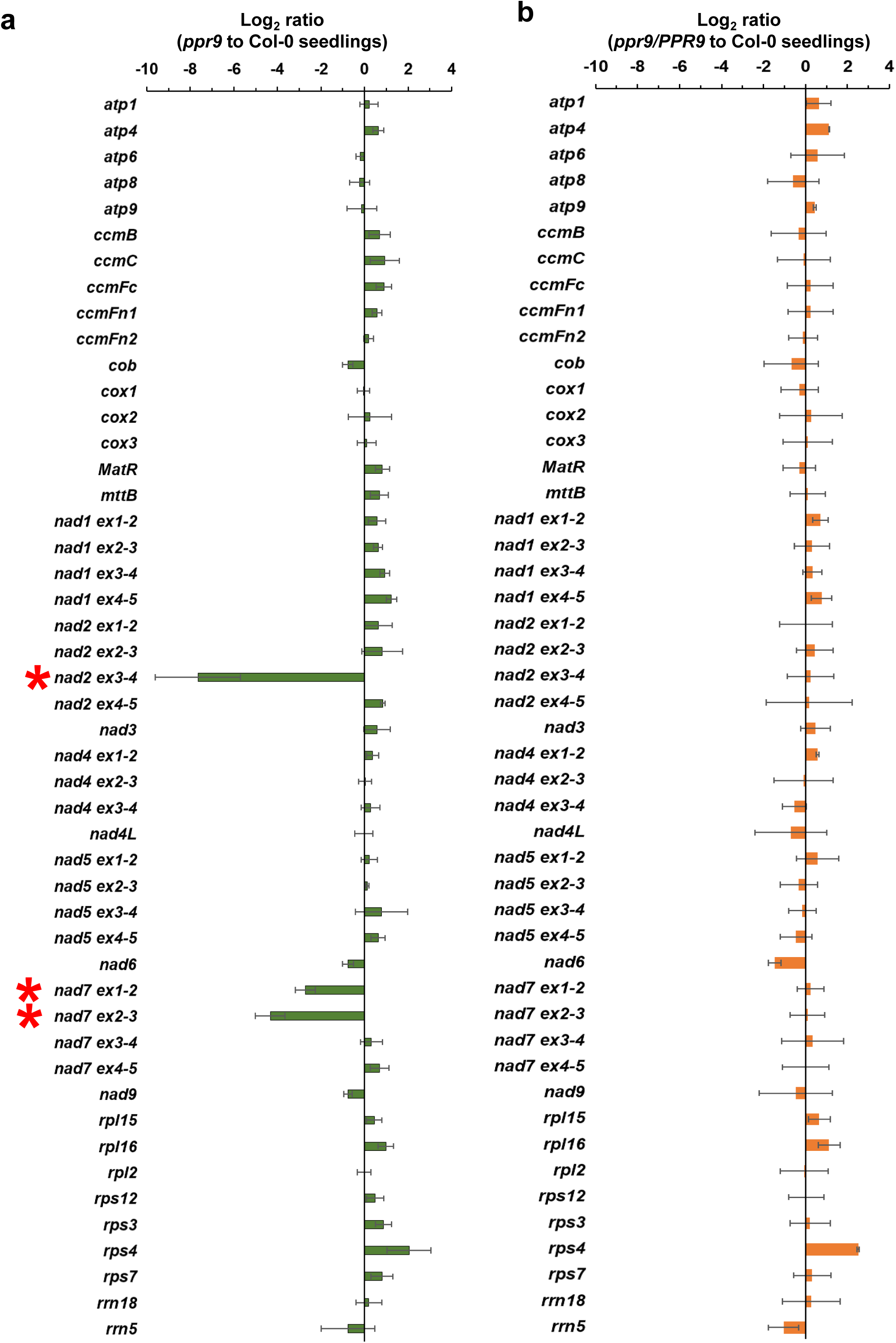
The accumulation of *nad2* and *nad7* transcripts relies on PPR9 in Arabidopsis mitochondria. The accumulation of various mitochondrial mRNA transcripts was analyzed by RT-qPCR. Total RNA isolated from 3-week-old Col-0 plants, 2∼3-month-old rescued *ppr9* mutant, *ppr9/PPR9* mutant plants, and plantlets obtained from Col-0 embryos at the torpedo stage, was reverse-transcribed, and the relative steady-state levels of cDNAs corresponding to the different organellar transcripts were estimated by qPCR with primers which specifically amplified specific organellar transcripts (Table S4). Histograms showing the relative mRNAs levels (i.e., log2 ratios) in rescued (a) homozygous *ppr9* or (b) *ppr9/PPR* plants, versus those of germinated embryos from wild type seeds. The values are means of nine RT-qPCRs corresponding to three biological replicates (error bars indicate one standard deviation), after normalization to the *GAPDH* (*AT1G13440*), *ACTIN2* (*AT3G18780*), *18S rRNA* (*AT3G41768*), and the mitochondrial *26S rRNA* (*rrn26*, *ATMG00020*) genes. Red asterisks indicate transcripts whose levels were significantly reduced in the mutant relative to the wild type control.

### PPR9 is required for the efficient splicing of *nad2* intron 3 and *nad7* introns 1 and 2

Based on the organellar mRNA profiles and reduced steady-state levels of *nad2* ex3-4 and *nad7* ex1-2 and ex2-3 transcripts in *ppr9* plantlets (Fig. 4), we considered defects in the splicing of *nad2* intron 3 and *nad7* introns 1 and 2, respectively. To test this assumption, we quantified the splicing efficiencies of the 23 mitochondrial introns in Arabidopsis mitochondria in germinated embryos of wild type plants, functionally complemented *ppr9/PPR9* plants and the *ppr9* mutant line. Splicing efficiency was evaluated by RT-qPCR as the ratio of specific pre-RNAs to their corresponding excised mRNAs in mutant lines relative to the wild type plantlets. An impairment in mitochondrial splicing was evident for *nad2* intron 3, with an approximately 1,000-fold reduction in splicing efficiency in *ppr9* relative to the wild type plants (Fig. 5a). The splicing of *nad7* introns 1 and 2 were also affected, showing about 30- and 60-fold reductions, respectively (Fig. 5a). While some Arabidopsis mutants defective in mitochondrial intron splicing also show increased accumulation of various other pre-RNAs, aside from defects in *nad2* intron 3 and *nad7* introns 1 and 2, the processing of other mitochondrial introns in *ppr9* was largely comparable to that of wild type plants (Fig. 5a). The processing of the organellar transcripts in *ppr9/PPR9* was equivalent to those of the germinated wild type embryos (Fig. 5b). These data indicate a specific role for PPR9 in the maturation of three mitochondrial group II-type introns in Arabidopsis plants, including *nad2* intron 3 and *nad7* introns 1 and 2.

**Figure 5.**
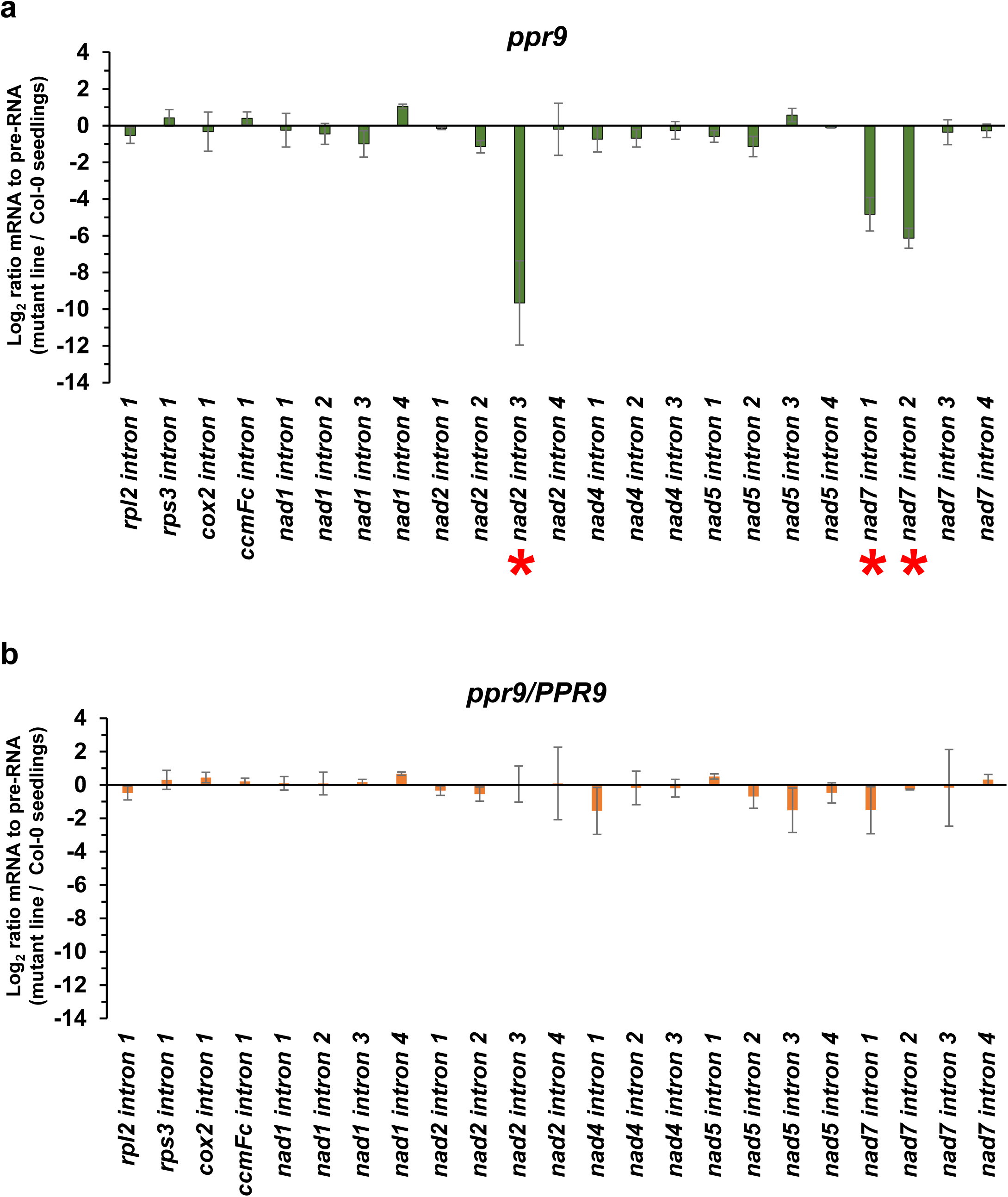
PPR9 is required for *nad2* intron 3 splicing and *nad7* transcript maturation in Arabidopsis mitochondria Total RNA was extracted from whole seedlings of wild type (Col-0), *ppr9/PPR9* and homozygous *ppr9* mutant plants. The relative (steady-state) levels of individual mitochondrial transcripts were evaluated by RT-qPCR, with oligonucleotides that designed to amplify specific organellar exon-exon or exon-intron regions (Table S4), using Col-0 RNA as the calibrator. For quantification, transcript levels were further normalized to various reference genes, including the *GAPDH* (*AT1G13440*), *ACTIN2* (*AT3G18780*), *18S rRNA* (*AT3G41768*), and the mitochondrial *26S rRNA* (*rrn26*, *ATMG00020*) genes. Splicing efficiencies of each of the 23 mitochondrial group II introns in *ppr9* (a) and functionally complemented *ppr9/PPR9* were assessed based on the relative pre-RNA to mRNA ratios in the mutant lines compared with those of wild type plants. The histograms show the log₂ fold change of mRNA/pre-RNA levels in the *ppr9* lines relative to Col-0. Data represent mean values from three biological replicates, each analyzed in triplicate (nine reactions in total). Error bars represent one standard deviation.

The RNA profiles were also analyzed by RNase-protection assays, which further indicated to altered processing of *nad2* intron 3 and *nad7* introns 1 and 2 (Fig. S4) in *ppr9*. The splicing efficiencies of the intron-containing *cox2* and *rpl2* transcripts, as well as the accumulation of an intronlerss (*nad6*) mRNA transcript, appeared to be unaffected (Fig. S4). We observed an accumulation of pre-RNA corresponding to the *trans*-spliced *nad2* intron 2 in the *ppr9* mutant in both the RNase-protection and RT-qPCR assays (Fig. S4). Since the processing of transcripts corresponding to matured *nad2* exon 2-3 RNAs seemed only slightly affected in the mutant (Figs. 4, 5, and S4), we speculate that an accumulation of precursor *nad2* intron 2 in *ppr9* likely results from the sequential nature of *nad2* splicing, where impaired processing of *nad2* intron 3 affects the processing of its related product, the precursor *nad2* intron 2. Accordingly, in multiple introns within a single gene (e.g., *nad1*, *nad2*, *nad4*, *nad5*, and *nad7*) the RNA processing steps seem to be interdependent. For example, the splicing or availability of one exon-intron segment may influence the substrate for the subsequent step, as in the cases of e.g., OTP43 [21] or MSP1 [3], where defects in *nad1* intron 1 processing also affect ‘downstream’ RNA processing steps.

### Analysis of respiratory complexes by native and denaturing PAGE

The respiratory system consists of five major protein complexes: CI (∼1,000 kDa), CII (∼160 kDa), a dimeric complex III (CIII₂, ∼500 kDa), CIV (200–220 kDa), and the ATP synthase enzyme (CV, ∼660 kDa). The incorporation of NAD2 and NAD7 subunits is essential for the assembly of a functional holo-complex I in plants [9, 55, 61, 69]. The reductions we see in the splicing efficiencies of several *nad2* and *nad7* pre-RNA transcripts (Figs. 4, 5) suggest that their corresponding subunits may be present at lower levels (or absent) in *ppr9* mutants. Accordingly, BN-PAGE analysis of Arabidopsis mitochondrial respiratory complexes revealed to altered CI biogenesis in homozygous *ppr9* mutants (Fig. 6). This was apparent by immunoblotting with antibodies against the carbonic anhydrase subunit CA2 (membranous domain) and the Nad9 protein (soluble arm) of CI. We further noticed the accumulation of various protein bands of lower molecular masses in the CA2 blots in *ppr9* plants. Similar CI-assembly intermediates of the membranous arm have been reported in Arabidopsis mutants defective in CI biogenesis [10, 49, 55, 61, 69]. We also compared the relative accumulation of mitochondrial complexes in germinated seedlings of wild type (Col-0) and the functionally complemented *ppr9/PPR9* line by BN-PAGE followed by immunoblotting with antibodies. These analyses also demonstrated that the levels of CI (with anti- CA2 and Nad9 antibodies), CIII (Rieske iron-sulfur protein, RISP), CIV (Cox2 subunit), and CV (AtpA/B subunits) in *ppr9/PPR9* seedlings were comparable to those observed in the wild type plants (Fig. 6 and Table S2a).

**Figure 6.**
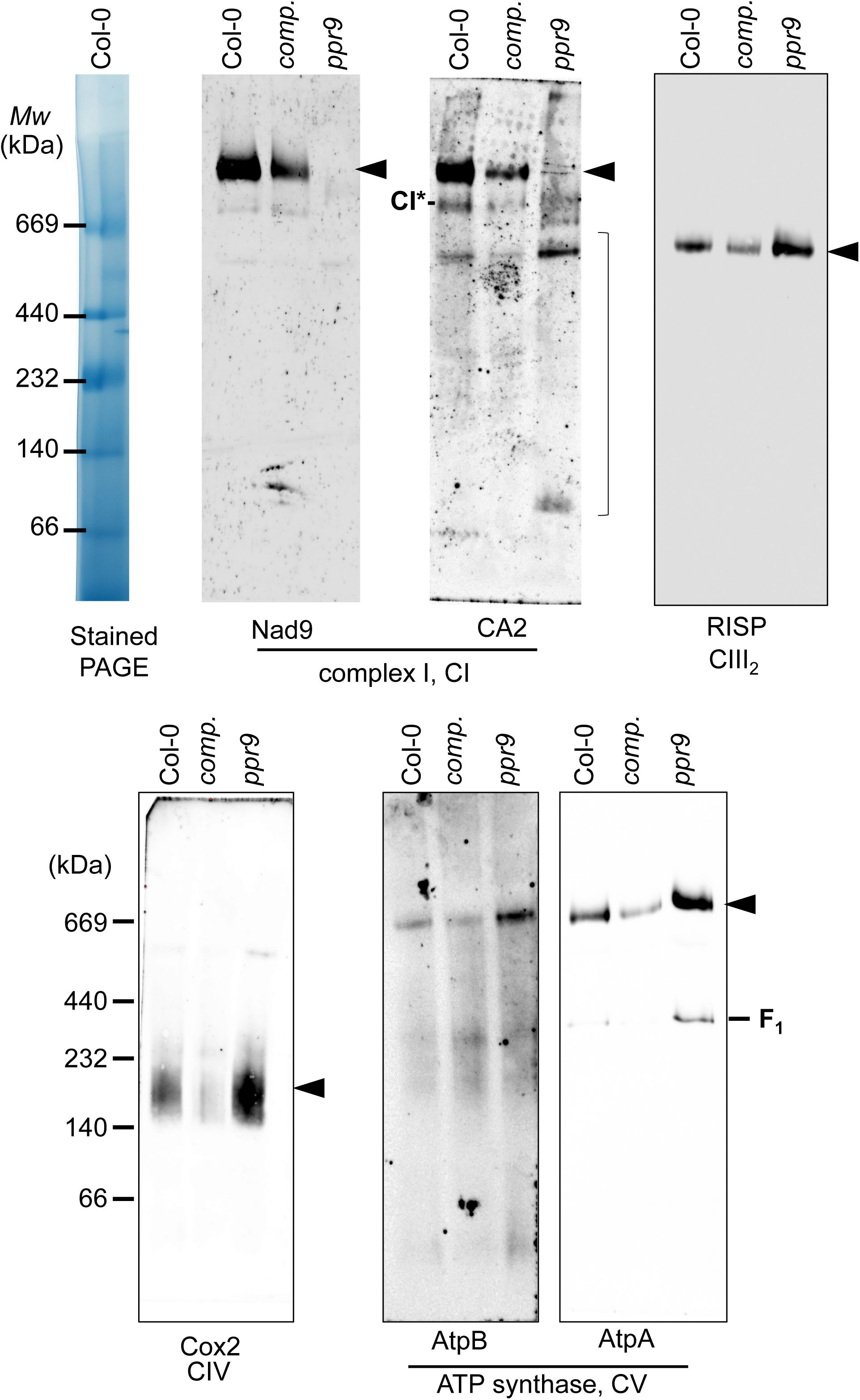
Defects in mitochondrial respiratory complex I biogenesis in *ppr9* mutants Accumulation of mitochondrial respiratory complexes in germinated embryos (at the torpedo stage) of wild type (Col-0), *ppr9/PPR9* and homozygous *ppr9* mutant plants. The abundance and assembly state of mitochondrial respiratory complexes were assessed by blue-native polyacrylamide gel electrophoresis (BN-PAGE) of crude organellar membrane preparations from Col-0 plants and ppr9 mutant lines. Following native electrophoresis, proteins were transferred to PVDF membranes and analyzed by immunoblotting using antibodies against various organellar protein subunits (Table S5), following chemiluminescence (ECL) assays and then recorded on an imaging system (ImageQuant LAS 4000, GE Healthcare, Haifa, Israel). Signal detection was performed by chemiluminescence. Arrows indicate the positions of the major native respiratory complexes, including NADH:ubiquinone oxidoreductase (CI; ∼1,000 kDa), cytochrome bc_1_ complex dimer (also called ubiquinol:cytochrome c oxidoreductase, or CIII₂; ∼500 kDa), cytochrome c oxidase (CIV; ∼220 kDa), and the ATP synthase enzymes (CV; ∼660 kDa). The asterisk in the CA2 panel marks the ∼850-kDa CI stable assembly intermediate (CI*), while additional lower-molecular-mass signals in the CA2 blot likely correspond to earlier CI-assembly intermediates [55, 69].

The steady-state levels of various mitochondrial proteins were also analyzed by denaturing SDS-PAGE followed by immunoblot analyses. The CA2 (∼30 kDa) subunit was found lower (about 50%, Table S2b) to those of the germinated wild type embryos (Col-0 heart) or functionally complemented *ppr9* line *(ppr9/PPR9*) (Fig. 7). The levels of NAD9 subunit (∼23 kDa) of the soluble arm of CI was more notably reduced (about 6% of that seen in the wild type plants) (Fig. 7, Table S2b). The signals of cytochrome b-c1 complex subunit Rieske-1 (RISP, ∼29 kDa) of complex III, cytochrome c oxidase subunit 2 (COX2, ∼30 kDa) and the Voltage-dependent anion-selective channel protein 1 (VDAC1, ∼29 kDa), were all found to be somewhat higher than those of the germinated wild type embryos collected from the same siliques (Fig. 7, Table S2b). Using equivalent amounts of fresh weight tissue, the respiratory complexes seemed to be similar in the functionally complemented *ppr9/PPR9* plants to those of the wild type plants (Fig. 7, Table S2b). In addition to the canonical respiratory complexes, plant mitochondria also harbor proteins of the alternative electron transport pathway, including various rotenone-insensitive NADH dehydrogenases (NDs) and alternative oxidases (AOXs) [9, 62, 65]. The steady-state level of AOX1a (∼33 kDa) was notably higher (∼250-fold) in *ppr9* mutant (Fig. 7, Table S2b).

**Figure 7.**
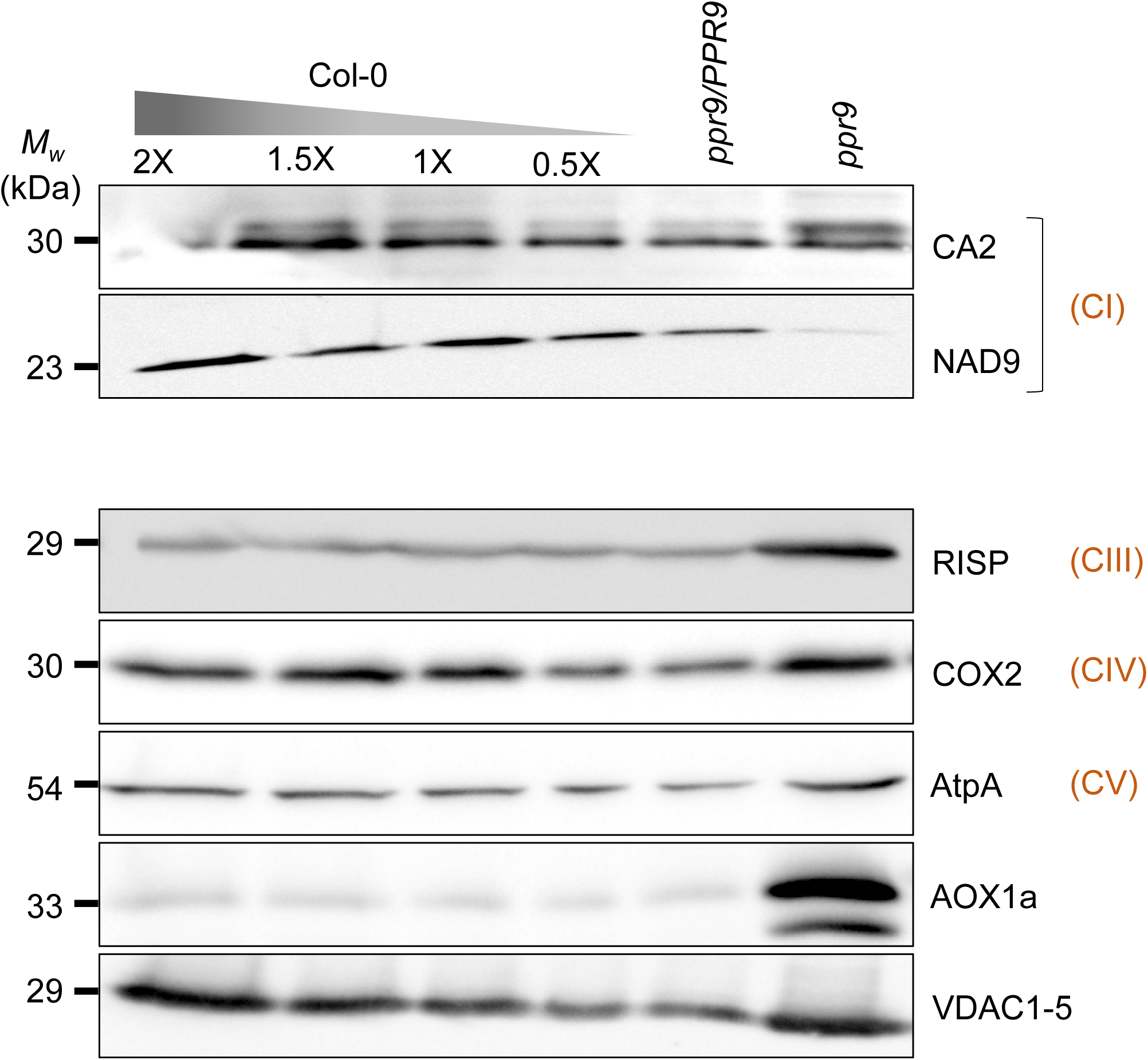
Accumulation of various organellar proteins in wild type and *ppr9* plants. Crude organellar (mitochondria-enriched) membrane fractions were isolated from wild type (Col-0), *ppr9/PPR9* and homozygous *ppr9* mutant plants. Equal amounts of protein were separated by SDS-PAGE and analyzed by immunoblotting with specific antibodies raised against representative organellar proteins (Table S5). Signals were detected using chemiluminescence (ECL) and recorded on an imaging system.

## Discussion

### *PPR9* encodes a mitochondrial P-type PPR protein essential for early embryogenesis

Mitochondria are central hubs of eukaryotic energy metabolism and hence are indispensable for plant physiology. Expression of mitochondrial genes in plants requires numerous post-transcriptional processing steps, including the splicing of group II introns embedded within the coding regions of many essential mitochondrial genes [17, 37, 92, 111]. These complex mt-RNA processing steps rely on a diverse set of RNA-binding proteins that may also serve as major regulatory cofactors for mitochondria biogenesis, respiratory activity, and consequently, plant growth and development. Here, we describe the molecular and biochemical analysis of PPR9, an essential P-type PPR protein factor in Arabidopsis encoded by the *AT1G03560* gene-locus.

Based on available databases (e.g., TAIR, UniProt, SUBA5) PPR9 encodes a lowly expressed mitochondria-targeted protein, harboring 13-to-14 PPR motifs (Figs. 1, S1). Publicly available MS/MS data [25] and our GFP localization analysis (Fig. 2) support the *in-silico* predictions and demonstrated that PPR9 reside in mitochondria *in vivo*. An embryo-rescue method (Figs. 3, S3) [88] allowed the establishment of homozygous *ppr9* (SALK_004994) mutant plants, which their RNA and protein profiles indicated that PPR9 is pivotal for mt-RNA processing and hence for mitochondria OXPHOS biogenesis (Figs. 4, 5, S4). Altered expression of Nad2 and Nad7 in *ppr9* mutants affects respiratory CI biogenesis (Figs. 6 and 7), which is strongly associated with defects in embryo development in the mutant Arabidopsis plants (Figs. 3 and S3).

### PPR9 promotes the splicing of several *nad2* and *nad7* group II-type introns

The mitochondrial introns in Arabidopsis mitochondria belong to the group II intron family [6, 11, 52, 83, 110]. Unlike canonical group II introns, whose splicing is facilitated by intron-encoded proteins (IEPs, or maturases) [52, 53, 64, 108], the splicing of mitochondrial introns in plants relies on a large and diverse set of nuclear-encoded protein cofactors. Some, such as PPR proteins, act specifically on a single intron target, whereas others, including Arabidopsis maturases, RNA helicases, and CRM- or PORR-related factors, function in the splicing of multiple introns [6, 11, 83, 92, 111]. PPR proteins constitute a particularly large family in angiosperms, comprising ∼450 members in Arabidopsis [2, 68, 84, 93]. These are postulated to bind single-stranded RNA in a modular and sequence-specific manner, and were shown to play key roles in organellar RNA metabolism, such as in RNA stabilization, editing, cleavage, and group II intron splicing [92, 94, 98, 111], and potentially also in mitoribosome biogenesis or translation [50, 78, 104]. Likewise, genetic characterization of *ppr9* mutants revealed a notable reduction in the steady-state levels of processed mitochondrial transcripts corresponding to *nad2* exons 3-4 and *nad7* exons 1-2 and exons 2-3 (Figs. 4, S4). RT-qPCR and RNase-protection assays further indicated that the removal of intron 3 of *nad2* and introns 1 and 2 of *nad7* was strongly affected in the mutant (Figs. 5, S4).

In some cases, *in-silico* PPR RNA-binding predictions [2, 15, 30, 86, 107] identify specific RNA-binding sites that correspond to experimentally validated biological targets. Yet, the current PPR code did not revealed a clear, specific binding site for PPR9. Instead, for PPR9 more than 200 putative binding sites were predicted across exonic, intronic and UTR sequences (excluding intergenic regions) (Table S1). These included *nad2* intron 3, and *nad7* introns 1 and 2 (Fig. 1B, Table S1), the three validated targets of PPR9, which appear to share common binding chrematistics. The molecular analysis of PPR9 suggests a refined binding code for PPR motifs that may improve the accuracy of identifying the *in vivo* targets of other PPR proteins. For example, the amino acids Met and Pro at positions 5 and 35, respectively, are predicted to bind A, U, or G; Arg and Asp may confer specificity for A; Asp and Pro are associated with binding to A or G ribonucleotides; and Thr and Met are linked to A, G, or U recognition. We also observed a single mismatch to the predicted consensus for the Thr–Thr combination at positions 5 and 35 of the 10^th^ PPR motif, where a C residue occurs instead of A or G (Fig. 1B).

### PPR9, holo-CI biogenesis, and embryo development

The mitochondrial electron transport chain (ETC) comprises of four main respiratory complexes (CI-CIV) that generate a proton gradient across the inner membrane, driving ATP biosynthesis by the ATP synthase enzyme (also denoted as CV) [9, 62, 65]. In addition to the canonical respiratory complexes, plants (and some other organisms) possess alternative oxidases (AOXs) and rotenone-insensitive NAD(P)H dehydrogenases (NDs) that can bypass the classical ETC system [9, 65]. CI, the largest respiratory complex (∼1,000 kDa), is composed of about 50 subunits, derived from expression of several mitochondrial gene-loci (*nad1*, *nad2*, *nad3*, *nad4*, *nad4L*, *nad5*, *nad6*, *nad7* and *nad9*) and numerous nuclear-encoded subunits that are imported into the organelle [45, 46, 58, 95]. These are assembled together in the organelles into two main subdomains (i.e., membrane-arm and the matrix-associated domain) that together form the functional holoenzyme, which is further assembled into respiratory chain supercomplexes (respirasomes) [9, 22, 28, 55, 61, 69, 85, 96]. The NAD2 and NAD7 subunits play pivotal roles in assembling the membrane and matrix domains, respectively [55]. Accordingly, the *ppr9* mutants that are defective in *nad2* and *nad7* RNA processing exhibited a notable reduction in holo-CI levels, as revealed by BN-PAGE (Fig. 6). Immunoblotting with the anti-CA2 antibodies further showed to distinct CI-related protein bands, including an ∼85 kDa particle (Fig. 6, CA2 panel), which was previously noted in *abo5* [55, 56] or *misf2* [67] mutants that are also affected in *nad2* expression. These particles, observed in *ppr9* and other mutants, likely represent CI assembly intermediates that are lacking NAD2.

Mutant plants defective in mitochondrial function, as well as those with impairments in cellular metabolism, such as fatty acid β-oxidation [72], display abnormal embryo development phenotypes, often with a ‘wrinkled’ seed morphology. Impaired CI biogenesis is likewise tightly associated with defective embryogenesis and subsequent growth and developmental abnormalities in Arabidopsis, where the degree of CI-deficiency has previously been correlated with phenotypic severity [17, 49, 69]. These abnormal growth and developmental phenotypes likely result from impaired cellular functions and reduced nutrient reserves in the seed. Likewise, homozygous *ppr9* mutants also exhibit early embryonic arrest, with *wrinkled* seed-morphology, failing to germinate under ‘standard’ growth conditions (Figs. 3a, S3).

In some cases, germination and seedlings establishment can be rescued under *in vitro* conditions, using immature seeds germinated on enriched growth media. The precise molecular mechanism underlying embryo- and plantlets growth and development rescue is still not fully understood. It is possible that the additional sugars, minerals and vitamins can support essential metabolic functions that compensate for metabolic constraints in mitochondrially affected mutant embryos. Also, prematurely harvested seeds with a softer seed coat may allow the emergence of the plantlets radicle and hypocotyl, by increasing permeability and reducing mechanical resistance [41]. The few germinated homozygous *ppr9* seedlings further displayed arrested growth and other phenotypes (Figs. 3c, S3), which are characteristic of CI-deficient plants [17, 49, 69]. While the external sugars and vitamins allowed limited vegetative development, the mutants could not complete their life cycle or produce viable seeds, underscoring the essentiality of PPR9 and mt-RNA maturation steps for mitochondria biogenesis, respiratory-related functions and plant physiology.

## Materials and methods

### Plant material and growth conditions

The experiments involved *Arabidopsis thaliana* (Col-0) plants. Seeds of the wild type and the *ppr9* mutants (SALK_004994) were obtained from the Arabidopsis Biological Resource Center (ABRC, Ohio State University, USA). For germination on half-strength (½) Murashige and Skoog (MS)-agar medium (M0221, Duchefa Biochemie, Haarlem, Netherland) wild type and mutant seeds were sterilized with Cl₂ gas and sown on MS-agar plates supplemented with 1% (w/v) sucrose or rescued using the method detailed below. After a 5-day stratification period at 4 °C, the plates were transferred to a growth chamber (Percival Scientific, Perry, IA, USA) set to 22°C with long-day conditions (16 h light/8 h dark), RH 50%, with a light intensity of 100∼150 µE m⁻² s⁻¹. The identity and integrity of each line were confirmed by PCR genotyping using gene-specific and T-DNA-specific primers (Table S3). Seeds from confirmed lines were sterilized, germinated on MS-agar medium containing 1% sucrose, and analyzed for their associated genetic and biochemical phenotypes. For larger-scale biomass production (e.g., for BN-PAGE analysis), seedlings from the MS-agar plates were transferred and grown in liquid MS medium supplemented with sucrose and vitamins (see below, ‘Embryo-rescue and establishment of a homozygous *ppr9* mutant line’), with moderate shaking (50–100 rpm) [88].

### Embryo-rescue and establishment of a homozygous *ppr9* mutant line

Embryo rescue of the *ppr9* mutant line was performed essentially as described previously [3, 88]. In brief, mature green siliques from heterozygous *ppr9* plants were surface-sterilized and then rinsed thoroughly with sterile DDW. Green (wild type or heterozygous) and white (homozygous *ppr9*) seeds isolated from the sterilized siliques were sown on MS-agar plates supplemented with 3% (w/v) sucrose and a vitamin mix (i.e., 100 μg myoinositol, 1.0 μg thiamine, 1.0 μg pyridoxine, 1.0 μg nicotinic acid per liter of media). To obtain larger quantities of plant material, required for BN-PAGE analyses, seedlings at the 6 leaves (L6) stage [8] were transferred to a liquid MS medium containing 1-3% sucrose and the same vitamin mix and grown under the conditions described previously, with gentle agitation (50-100 RPM), as we performed previously [3, 88].

### Imaging-based analysis of Arabidopsis plant tissues

For morphological analyses, plant tissues were examined using a stereoscopic (dissecting) microscope or a light microscope at the Bio-Imaging Unit, Institute of Life Sciences, The Hebrew University of Jerusalem.

### Protoplasts preparation and GFP localization assays

GFP fusion proteins were expressed in protoplasts prepared from *Physcomitrium patens* [100, 101, 105], and transformed generally as described previously [13]. In brief, protoplasts were obtained from harvested tissue of a 1-month-old liquid culture (protonemata enriched) and treated with 4% (w/v) Driselase in 0.5 M mannitol. The resulting suspension was filtered through a 45 µm sieve. A DNA product encoding the first 82 amino acids of the intronless *PPR9* gene, fused in-frame to eGFP, was cloned into the binary pCAMBIA-1302 vector (CAMBIA, Canberra, Australia; https://www.cambia.org; GenBank AF234298.1) under the control of CaMV 35S promoter and a NOS terminator. The pCAMBIA-1302-N-PPR9-eGFP construct was then introduced into the *P. patens* protoplasts by PEG 4000 transformation method [80]. For transformation, protoplasts were pelleted by centrifugation (10 min, 50 ×g, at 25 °C), and approximately 1.2 × 10⁵ cells were incubated with 30∼50 µg of plasmid DNA suspended in 0.1 M Ca(NO_3_)_2_ within a 25% (w/v) PEG 4000 solution. Transformed cells were imaged 72 h post-transformation using an Olympus FV3000 confocal microscope, and the resulting images were visualized and processed with the Fiji open-source software [82].

### Mitochondrial transcript analysis in wild type and mutant plants

Total RNA was extracted from approximately 50 mg of plant tissue using RNAzol® RT (R4533, Sigma-Aldrich, Rehovot, Israel) reagent. The isolated RNA was subsequently treated with RNase-free DNase I to remove DNA prior for its use in RT-qPCR analysis. In brief, cDNA was synthesized from 1-3 µg of total RNA using SuperScript III First-Strand Synthesis System for RT-PCR (Cat. no. 18080-051, Invitrogen, Rhenium Modi’in, Israel). RNA (11 µl of total RNA in nuclease-free water) was mixed with 1 µl dNTP mix (2.5 mM each nucleotide) and 1 µl random hexamers (1 µg/ml), incubated at 65 °C for 5 min, and immediately chilled on ice for 5 min. A reaction mix containing 4 µl 5× ‘First Strand’ buffer, 1 µl DTT, 1 µl RNasin RNase inhibitor, and 1 µl SuperScript III, for each cDNA synthesis reaction, was prepared on ice, and added to each RNA sample. Samples were then incubated at 25°C for 5 min for primer annealing, followed by 60 min at 50°C for cDNA synthesis, and 15 min at 70°C to inactivate the enzyme. Gene-specific oligonucleotides were designed to amplify both mRNA and pre-mRNA transcripts (Table S4) in qPCR. For normalization, *Arabidopsis thaliana* reference genes, *GAPDH* (*AT1G13440*), *ACTIN2* (*AT3G1878*), and *18S rRNA* (*AT3G41768*), were used as internal calibrators. RNase-protection assays [16, 43, 44, 70] were performed with *in vitro* transcribed labeled RNA fragments complementary to exon-intron regions in each transcript. DNA fragments, amplified by PCR using a 5′ primer containing the T7 RNA polymerase promoter sequence fused to the gene of interest, were used to generate RNA probes (labeled with Fluorescein-12-UTP, Cat. 11427857910, Sigma-Aldrich Israel Ltd, Rehovot, Israel) complementary to organellar transcripts. Detection was carried out using the Molecular Dynamics Storm system, and analyzed by the ImageQuant software at the Center for Molecular Dynamics (HUJI, Jerusalem, Israel).

### Enrichment of organellar membranes from wild type and mutant seedlings

Crude organellar membrane fractions from wild type and mutant plants were prepared as previously described [71] with minor adjustments [89]. Approximately 200 mg of whole plant tissue was homogenized in 2.0 mL of membrane extraction buffer (75 mM MOPS-KOH, pH 7.6; 0.6 M sucrose; 4 mM EDTA; 0.2% polyvinylpyrrolidone-40; 8 mM L-cysteine; and 0.2% bovine serum albumin). The homogenate was filtered through a single layer of Miracloth to remove cell debris and centrifuged at 1,300 × g to pellet cell fragments and thylakoid membranes. The crude mitochondrial fraction was collected by a sequential centrifugation at 22,000 ×g for 10 minutes at 4°C.

### Analysis of the steady state levels of organellar proteins in wild type and mutant plants

Protein was extracted from crude organellar membrane fractions corresponding to approximately 200 mg of fresh whole-plant tissue, solubilized in protein sample buffer, and separated by SDS-PAGE [51]. For immunoblotting, the proteins were transferred onto a PVDF membrane and incubated overnight at 4°C with specific antibodies (Table S5). Detection was performed using chemiluminescence following incubation with horseradish peroxidase (HRP)-conjugated secondary antibody.

### Blue Native (BN-) PAGE analysis of mitochondrial respiratory complexes

BN-PAGE of crude organellar membranous fractions was performed generally according to the method described by Pineau, et al. [71]. In brief, crude organellar membrane fraction, equivalent of 200 mg MS-agar grown Arabidopsis seedlings, was solubilized in 200 μL of 1.5% (w/v) n-Dodecyl β-D-maltoside (DDM, CAS no. 69227-93-6) solution containing 0.6% (w/v) Brilliant Blue G (CAS no. 6104-58-1). Insoluble materials were removed by a centrifugation step (i.e., 20,000 xg). Solubilized membrane fractions were loaded onto a native gel. For immunoblotting, the proteins were transferred to a PVDF membrane, and incubated with primary antibodies (Table S5), as indicated in each blot. Detection was carried out by chemiluminescence assay with horseradish peroxidase (HRP)-conjugated secondary antibody.

### Estimation of the relative accumulation of different organellar proteins in wild type (Col-0) and *ppr9* mutant plants

Detection of organellar proteins was performed by immunoblotting assays and using the ImageQuant™ LAS-4000 system (GE Healthcare Life Sciences, Daniel Biotech, Rehovot, Israel) following chemiluminescence assays with horseradish peroxidase (HRP)-conjugated secondary antibodies. The signals were analyzed and quantified using the Fiji open-source software [82].

### Data statement

The Arabidopsis Information Resource (TAIR) database provides genetic and molecular biology data for the model plant *Arabidopsis thaliana* [74]. The predicted three-dimensional structures of proteins were obtained from and analyzed using the AlphaFold Protein Structure Database, which provides publicly accessible computational models of protein structures [103]. Accession numbers: Arabidopsis mitochondrial genome, BK010421.1; PPR9: gene AT1G03560, protein Q9LR67 (PPR9_ARATH). Data supporting the results of this study can be obtained from the corresponding author upon reasonable request.

## Acknowledgements

The authors thank Dr. Yoram Soroka and Dr. Haim Azulay (The Hebrew University of Jerusalem, HUJI) for their assistance with mutant analyses. We also thank Prof. Dr. Ute Höcker (University of Cologne, Germany) for providing *P. patens* lines and protocols for protoplasts preparation and transformation. Confocal microscopy was assisted by Dr. Yang-Sung Sohn (HUJI) and supported by the ‘Charles E. Smith Family’ and the ‘Prof. Joel Elkes Laboratory for Collaborative Research in Psychobiology’ (HUJI). This work was supported by grants to O.O.B. from the Israel Science Foundation (ISF-911/25).

## Short legends for Supporting Information

**Table S1.** PPR9 mt-RNA gene-loci binding predictions.

**Table S2**. Quantification data of protein profiles.

**Table S3.** List of oligonucleotides used for the screening of *ppr9* mutants in this study.

**Table S4.** List of RT-qPCR oligonucleotides designed to specific different exon-exon (analysis of mRNA and processed transcripts) and intron-exon regions (analysis of pre-RNAs) in Arabidopsis mitochondria.

**Table S5.** List of antibodies used for the analysis of organellar proteins in wild type and *ppr9* mutant plants.

**Figure S1.** PPR9 protein structure and its predicated RNA biding site.

**Figure S2**. Expression patterns of PPR9 across tissues and developmental stages.

**Figure S3**. Developmental phenotypes associated with embryo-rescued *ppr9* mutants.

**Figure S4**. mt-RNA metabolism defects associated with embryo-rescued *ppr9* mutants.

## Author contributions statement

E.K., and M.K., Plant growth and analysis, establishment of embryo rescued mutant lines, DNA, RNA and protein analyses. R.Ma, GFP localization analyses. R.Mi., Help with protein folding analysis and RNase-protection assays. N.C., *in-silico* predictions of PPR9 RNA targets. O.O.B., PI, manuscript preparation and corresponding author.

