## Supplementary Data for "PPR9 mediates mitochondrial *nad* transcript maturation required for complex I biogenesis and early plant development in Arabidopsis" (from another version of this preprint: version 1, 2025-12-05)

**Supplementary Data Files**

**Supplementary Tables:**

**Table S1.** PPR9 mt-RNA gene-loci binding predictions.

**Table S2**. Quantification data of protein profiles.

**Table S3.** List of oligonucleotides used for the screening of *ppr9* mutants in this study.

**Table S4.** List of RT-qPCR oligonucleotides designed to specific different exon-exon (analysis of mRNA and processed transcripts) and intron-exon regions (analysis of pre-RNAs) in Arabidopsis mitochondria.

**Table S5.** List of antibodies used for the analysis of organellar proteins in wild type and *ppr9* mutant plants.

**Supplementary Figure legends**

**Figure S1.** PPR9 protein structure and its predicated RNA biding site.

(a) The deduced amino acid sequence of PPR9 (UniProt Q9LR67 · PPR9_ARATH). The predicted mitochondrial targeting sequence (53 amino acid long) is highlighted in magenta, with the cleavage sites indicated by arrow and line. Regions of low confidence are colored in black. Predicted PPR motifs are coded blue for regions of high confident, whereas domains of low scoring are highlighted in cyan. (B) Homology modeling of *Arabidopsis thaliana* (Col-0) PPR9. The 3D model of the PPR9 protein structure was obtained from AlphaFold server (Jumper et al., 2021). The hypothetical 3D surface was visualized with PyMol. The “i” to “iii” panels represent ribbon structures at different angles, whereas “iv” to “vi” indicate the hypothetical surface of PPR9. The color code is red for negative values, white for near zero values and blue for positive values.

**Figure S2**. Expression patterns of PPR9 across tissues and developmental stages.

Expression levels and patterns of PPR9 were analyzed using the publicly available ‘The Arabidopsis Information Resource’ website (TAIR; http://www.arabidopsis.org).

**Figure S3**. Developmental phenotypes associated with embryo-rescued *ppr9* mutants.

Siliques at the green-mature stage were collected from heterozygous *ppr9* plants. These siliques contain typical green seeds (wild-type or heterozygous) and translucent/pale-green seeds that harbor homozygous *ppr9* mutant embryos at the late torpedo stage. Green and translucent/pale-green seeds obtained from surface-sterilized siliques were germinated and grown on MS agar plates supplemented with 3% sucrose until they reached the L6 stage (Boyes *et al.* 2001), and were subsequently transferred to liquid medium for further biomass production (Best *et al.* 2023, Shevtsov-Tal *et al.* 2021). The image shows *in vitro*-germinated wild-type (Col-0), functionally complemented *ppr9/PPR9* plants and eight individual ~2-month-old homozygous *ppr9* seedlings grown in liquid medium at 22°C and 50% relative humidity, under long-day conditions (18 h light / 6 h dark) with a light intensity of ~100 µE $m^{-2}$ $s^{-1}$. The upper panel shows Col-0, *ppr9/PPR9*, and homozygous *ppr9* plantlets. The middle panel illustrates a phenotypic variation typical of rescued *ppr9* mutants (i.e., plantlets *ppr9-2* to *ppr9-4*), with several plantlets exhibiting a bushy-like morphology. Many of the germinated seedlings (e.g., *ppr9-5* and *ppr9-6*) failed to develop and died shortly after being transferred from MS plates to liquid medium, whereas others (e.g., plantlets *ppr9-7* and *ppr9-8*) collapsed after several weeks in liquid culture. Genotypes were confirmed by PCR and sequencing. Scale bars represent 1 cm, as indicated in each panel.

**Figure S4**. mt-RNA metabolism defects associated with embryo-rescued *ppr9* mutants.
Total RNA was extracted from wild-type (Col-0) and homozygous *ppr9* mutant plants. The relative (steady-state) accumulation of specific mitochondrial transcripts was evaluated by RNase protection assays and RT-qPCR analyses, using probes or oligonucleotides (respectively) (Table S4) designed to specific exon-exon or exon-intron regions, with Col-0 RNA used as calibrator. The histograms show the relative accumulation (i.e., $Log_2$ ratios) of mRNA (light blue) and pre-RNA (orange) transcripts in *ppr9* mutants compared with those of germinated embryos of Col-0 plants. Values represent the means of nine RT-qPCRs corresponding to 3 biological replicates (error bars represent one standard deviation), normalized to *GAPDH* (*AT1G13440*), *ACTIN2* (*AT3G18780*), *18S rRNA* (*AT3G41768*), *RRN26* (*mitochondrial 26S rRNA*, *ATMG00020*), *RRN5* (*ATMG01380*), and *RRN18* (*ATMG01390*). Schematic gene structures are shown below the representative RNA blots, with scale bar represents 500 bp.

**Table S1**. PPR9 mt-RNA gene-loci binding predictions.

a. PPR motif / RNA-base binding prediction.

| **PPR motif** | **Amino acids at 5' and 35'** | **related RNA base** |
|---|---|---|
| (PPR-0)* | - | - |
| PPR-1 | NT | C>U |
| PPR-2 | ND | U>C>G |
| PPR-3 | ND | U>C>G |
| PPR-4 | MP | N |
| PPR-5 | SN | A |
| PPR-6 | TD | G>>A |
| PPR-7 | SN | A |
| PPR-8 | SD | G>>A>U |
| PPR-9 | NT | C>U |
| PPR-10 | TT | A>G |
| PPR-11 | RD | N |
| PPR-12 | DP | N |
| PPR-13 | TM | N |

* - a degenerated domain

b. mt-RNA gene-loci related to the predicted binding site of PPR9.

| **Target** | **mtDNA loci** | | **Gene** | **Feature** | **Strand** |
|---|---|---|---|---|---|
| | **Start** | **End** | | | |
| CTTTAGAATGAAA | 96692 | 96704 | *nad2 intron 3 (nad2.2)* | intron | - |
| TGTAAGAGCCAGG | 244582 | 244570 | *nad7 intron 1* | intron | - |
| CTGGAAATTGAAT | 243506 | 243518 | *nad7 intron 2* | intron | - |
| Additional gene-loci predicted for the PPR9 binding code, ranked according to their matching scores | | | | | |
| CTTGAGAATGAGA | 68408 | 68420 | *atp1* | exon | - |
| TCTGAGAGTGAGA | 68021 | 68033 | *atp1* | exon | - |
| TTGAAAAACAAAT | 67288 | 67300 | *atp1* | exon | - |
| TGGAAAAACGACT | 68085 | 68097 | *atp1* | exon | - |
| TTCTAAAGCGGTA | 68655 | 68667 | *atp1 5'UTR* | 5'UTR | - |
| CTCGAAAGCAGAA | 362706 | 362718 | *atp4* | exon | + |
| CTTAAGAATGGTT | 362751 | 362763 | *atp4* | exon | + |
| TTTCAAAGTGACT | 362373 | 362385 | *atp4* | exon | + |
| TTTGAGATTGTCC | 49837 | 49849 | *atp6-1* | exon | + |
| TTTGAGATTGTCC | 265198 | 265210 | *atp6-2* | exon | - |
| TTGAAAAATATCA | 265269 | 265281 | *atp6-2* | exon | - |
| TGCAAAATCAATA | 271162 | 271174 | *atp9* | exon | - |
| CCCGAAAATGCCC | 270804 | 270816 | *atp9 3'UTR* | 3'UTR | - |
| TCCAAGATCAAAG | 271268 | 271280 | *atp9 5'UTR* | 5'UTR | - |

| | | | | | |
|---|---|---|---|---|---|
| TCCGAGAATGTTG | 17763 | 17775 | *ccmB 3'UTR* | 3'UTR | - |
| TTCCAGAACAGGA | 191629 | 191641 | *ccmFC exon 2* | exon | + |
| TCTTAAAACAAGG | 316335 | 316347 | *ccmFn1* | exon | + |
| CCGTAGAACGTAT | 292086 | 292098 | *ccmFn2* | exon | + |
| CGGTAAATCGGGG | 291544 | 291556 | *ccmFn2 5'UTR* | 5'UTR | + |
| TTTAAAAGTATGT | 181907 | 181919 | *cob* | exon | - |
| CCTTAAATCGTTT | 182305 | 182317 | *cob* | exon | - |
| TTCCAGATTATCC | 115009 | 115021 | *cox1* | exon | - |
| TTTAAGAGTAAGG | 8397 | 8409 | *cox2 3'UTR* | 3'UTR | + |
| TTCTAAAATGGTT | 6271 | 6283 | *cox2 exon 1* | exon | + |
| TCCAAGATCGTAC | 7463 | 7475 | *cox2 intron* | intron | + |
| TTTGAGATTGTTC | 329737 | 329749 | *cox3 3'UTR* | 3'UTR | - |
| CCCGAAATTAGAT | 329759 | 329771 | *cox3 3'UTR* | 3'UTR | - |
| TGGCAGATCATGT | 329786 | 329798 | *cox3 3'UTR* | 3'UTR | - |
| TCTCAAAGTAATA | 330935 | 330947 | *cox3 5'UTR* | 5'UTR | - |
| TTGTAAATCAAGA | 232941 | 232953 | *matR* | exon | + |
| TGGAAAAGCGTCT | 232215 | 232227 | *matR* | exon | + |
| TCGTAAATTGGTC | 232758 | 232770 | *matR* | exon | + |
| TGGCAAAGTAACT | 218912 | 218924 | *mttB* | exon | - |
| TGCCAAATCGTCG | 219068 | 219080 | *mttB* | exon | - |
| CCCGAAAATGCCC | 234186 | 234198 | *nad1 3'UTR (nad1.3 )* | 3'UTR | + |
| TTCAAGAATGTCT | 83850 | 83862 | *nad1 5'UTR (nad1.1)* | 5'UTR | - |
| TTTTAGAATGGCT | 83091 | 83103 | *nad1 exon 1 (nad1.1)* | exon | - |
| TTCGAAATTGATT | 83145 | 83157 | *nad1 exon 1 (nad1.1)* | exon | - |
| CGCAAAAGCAGAT | 59339 | 59351 | *nad1 exon 3 (nad1.2)* | exon | + |
| TGGGAGAGTATGC | 230332 | 230344 | *nad1 exon 4 (nad1.3 )* | exon | + |
| CCGGAAAGTGTTC | 234051 | 234063 | *nad1 exon 5 (nad1.3 )* | exon | + |
| TTTGAGAACATAT | 82267 | 82279 | *nad1 intron 1.1 (nad1.1)* | intron (trans splicing) | - |
| CCGTAAATTGAGG | 82571 | 82583 | *nad1 intron 1.1 (nad1.1)* | intron (trans splicing) | - |
| CCTGAAATTACAT | 57583 | 57595 | *nad1 intron 1.2 (nad1.2)* | intron (trans splicing) | + |
| CGTGAGAGCAAGG | 58688 | 58700 | *nad1 intron 2 (nad1.2)* | intron | + |
| TCTAAAAGCGGGA | 58826 | 58838 | *nad1 intron 2 (nad1.2)* | intron | + |
| TCGGAAAGCACGG | 59220 | 59232 | *nad1 intron 2 (nad1.2)* | intron | + |
| TGCGAGAATAACT | 58788 | 58800 | *nad1 intron 2 (nad1.2)* | intron | + |
| TCTCAAAGTAAAG | 233459 | 233471 | *nad1 intron 4 (nad1.3 )* | intron | + |
| CGCGAGAGTGTCA | 233779 | 233791 | *nad1 intron 4 (nad1.3 )* | intron | + |
| TCTAAGATTAGGG | 230547 | 230559 | *nad1 intron 4 (nad1.3 )* | intron | + |
| TTCAAGATCACTG | 97894 | 97906 | *nad2 exon 3 (nad2.2)* | exon | - |
| TTTTAGAATGTTG | 163396 | 163408 | *nad2 intron 2.1 (nad2.1)* | intron (trans splicing) | + |
| CGCGAAATCAGAG | 163633 | 163645 | *nad2 intron 2.1 (nad2.1)* | intron (trans splicing) | + |
| CCGCAAAGCAAAA | 97067 | 97079 | *nad2 intron 3 (nad2.2)* | intron | - |
| CTTAAAATTATGT | 95972 | 95984 | *nad2 intron 3 (nad2.2)* | intron | - |
| TGCTAAAGCACCT | 97170 | 97182 | *nad2 intron 3 (nad2.2)* | intron | - |
| CGGGAAAGCGCTG | 95618 | 95630 | *nad2 intron 3 (nad2.2)* | intron | - |
| TCCTAGATTGAAG | 96880 | 96892 | *nad2 intron 3 (nad2.2)* | intron | - |

| | | | | | |
|---|---|---|---|---|---|
| TCGGAGATCGAGA | 95830 | 95842 | *nad2 intron 3 (nad2.2)* | intron | - |
| CCCCAAAGTAGAA | 94320 | 94332 | *nad2 intron 4 (nad2.2)* | intron | - |
| TTCAAGAATACGA | 215458 | 215470 | *nad4 exon 1* | exon | - |
| CCCGAAAGCGTGC | 215115 | 215127 | *nad4 exon 1* | exon | - |
| TGTTAGAACATTT | 215549 | 215561 | *nad4 exon 1* | exon | - |
| CGTCAAAGTGCCT | 213404 | 213416 | *nad4 exon 2* | exon | - |
| CCCCAAAGTGTTC | 207637 | 207649 | *nad4 exon 4* | exon | - |
| TCTCAGAATAAAA | 214306 | 214318 | *nad4 intron 1* | intron | - |
| TTCAAGAATAGGG | 214334 | 214346 | *nad4 intron 1* | intron | - |
| CCGAAAAGTAAAA | 213823 | 213835 | *nad4 intron 1* | intron | - |
| CGCGAGAATGGAT | 214754 | 214766 | *nad4 intron 1* | intron | - |
| TGCTAAAACAAAG | 214471 | 214483 | *nad4 intron 1* | intron | - |
| TCGTAAAGTAGTC | 214997 | 215009 | *nad4 intron 1* | intron | - |
| CGGCAAATCAAAA | 214380 | 214392 | *nad4 intron 1* | intron | - |
| TTGAAGAGTACCT | 210323 | 210335 | *nad4 intron 2* | intron | - |
| TCGGAAAGCACCC | 211066 | 211078 | *nad4 intron 2* | intron | - |
| TTTCAAATCGGAT | 212702 | 212714 | *nad4 intron 2* | intron | - |
| CCGCAGAACGAGA | 210953 | 210965 | *nad4 intron 2* | intron | - |
| CCGTAGATCGGAA | 211073 | 211085 | *nad4 intron 2* | intron | - |
| TCGTAAATCAAAA | 212822 | 212834 | *nad4 intron 2* | intron | - |
| TCGGAGAGCACTT | 207733 | 207745 | *nad4 intron 3* | intron | - |
| TGGCAGAGCACGA | 209160 | 209172 | *nad4 intron 3* | intron | - |
| CTCGAAATTAAAT | 234338 | 234350 | *nad5 5'UTR (nad5.1)* | 5'UTR | + |
| TTTCAAAGCATTA | 236211 | 236223 | *nad5 exon 2 (nad5.1)* | exon | + |
| CTTCAAAGTGGAT | 27194 | 27206 | *nad5 exon 4 (nad5.3)* | exon | + |
| TCGGAGAGCACTG | 235304 | 235316 | *nad5 intron 1 (nad5.1)* | intron | + |
| CGGAAGATCAAGT | 235255 | 235267 | *nad5 intron 1 (nad5.1)* | intron | + |
| CTTGAGAATAAAG | 360217 | 360229 | *nad5 intron 3.1 (nad5.2)* | intron (trans splicing) | + |
| CCCAAGAGCGAGG | 360248 | 360260 | *nad5 intron 3.1 (nad5.2)* | intron (trans splicing) | + |
| CTGAAAAGCAGTT | 27515 | 27527 | *nad5 intron 4 (nad5.3)* | intron | + |
| TGCTAGAGCACGG | 27918 | 27930 | *nad5 intron 4 (nad5.3)* | intron | + |
| CGGGAGAATGAAG | 27928 | 27940 | *nad5 intron 4 (nad5.3)* | intron | + |
| TTGTAAAGTAGTA | 245496 | 245508 | *nad7 5'UTR* | 5'UTR | - |
| TGGAAAAACAAAA | 245564 | 245576 | *nad7 5'UTR* | 5'UTR | - |
| CCCAAGAACACGC | 243062 | 243074 | *nad7 exon 3* | exon | - |
| TTTCAAATCATTC | 242960 | 242972 | *nad7 exon 3* | exon | - |
| TCGTAAAATAAGA | 239367 | 239379 | *nad7 exon 5* | exon | - |
| CCCAAGAACAACC | 241920 | 241932 | *nad7 intron 3* | intron | - |
| TCTGAGAATGATT | 241701 | 241713 | *nad7 intron 3* | intron | - |
| TCCAAGATCAGAA | 240889 | 240901 | *nad7 intron 4* | intron | - |
| TTGGAAAGTACCC | 240272 | 240284 | *nad7 intron 4* | intron | - |
| TGGGAAAACGCTG | 240373 | 240385 | *nad7 intron 4* | intron | - |
| CTCGAAAACGAAG | 24861 | 24873 | *nad9* | exon | + |
| TCTCAAATTGCAT | 24796 | 24808 | *nad9* | exon | + |
| TTGAAGAGCAAGA | 24620 | 24632 | *nad9 5'UTR* | 5'UTR | + |

| | | | | | |
|---|---|---|---|---|---|
| TCCAAAATCATAA | 328398 | 328410 | *ORF100A* | ORF | + |
| CCTCAGAATAAGC | 226458 | 226470 | *ORF102B* | ORF | - |
| TTCCAGAGTACCT | 257724 | 257736 | *ORF106B* | ORF | + |
| CTTGAAAGCGGTG | 309370 | 309382 | *ORF106D* | ORF | + |
| CTCGAAAGCAAAG | 284716 | 284728 | *ORF106E* | ORF | + |
| TCGGAAAACAAAC | 275131 | 275143 | *ORF107G* | ORF | - |
| TCCCAAAGCACTA | 327198 | 327210 | *ORF109B* | ORF | + |
| TGGTAGAGCAGTT | 164997 | 165009 | *ORF110A* | ORF | + |
| CTGAAAAATGGCT | 367643 | 367655 | *ORF110B* | ORF | - |
| CGGTAGAATGTGG | 367513 | 367525 | *ORF110B* | ORF | - |
| TTCTAAAGTGCGA | 70411 | 70423 | *ORF113* | ORF | - |
| TGGTAGAGTACAA | 70383 | 70395 | *ORF113* | ORF | - |
| TTCTAAAGTACCA | 13010 | 13022 | *ORF116* | ORF | - |
| TTTCAAAGTATAT | 290770 | 290782 | *ORF117* | ORF | + |
| TCTAAAATCAAGA | 290792 | 290804 | *ORF117* | ORF | + |
| TTCGAGATCAAAG | 287700 | 287712 | *ORF118* | ORF | + |
| TTTAAAAATGGGT | 327888 | 327900 | *ORF119* | ORF | - |
| CGGTAGATTATGT | 327900 | 327912 | *ORF119* | ORF | - |
| CTTGAGAACGGAT | 249797 | 249809 | *ORF122A* | ORF | + |
| CGGGAAAGCACTT | 90540 | 90552 | *ORF136A* | ORF | + |
| CTCTAGAACAGTT | 91306 | 91318 | *ORF136B* | ORF | + |
| TTTGAGATTGAGA | 91208 | 91220 | *ORF136B* | ORF | + |
| CCCCAAAATGCCA | 438 | 450 | *ORF139A* | ORF | - |
| CCCCAAAATGCCA | 194635 | 194647 | *ORF139B* | ORF | - |
| CGGCAGAACATGT | 246418 | 246430 | *ORF141* | ORF | - |
| CTCCAAAACAGCC | 16531 | 16543 | *ORF143* | ORF | + |
| CGCGAGAGTAGGG | 74119 | 74131 | *ORF145B* | ORF | + |
| CGGGAAATCGGTT | 54393 | 54405 | *ORF152A* | ORF | + |
| CGGGAAATCGGTT | 260642 | 260654 | *ORF152B* | ORF | - |
| TGCAAGAGTATTC | 277075 | 277087 | *ORF159* | ORF | + |
| TGGAAGAACACCA | 225691 | 225703 | *ORF160* | ORF | + |
| CTGAAGAGCGCTC | 311623 | 311635 | *ORF187* | ORF | - |
| TTTTAAATTATTC | 42786 | 42798 | *ORF199* | ORF | - |
| CCCAAGAGCAGCA | 346950 | 346962 | *ORF240A* | ORF | + |
| TTCGAGAACAAAT | 346630 | 346642 | *ORF240A* | ORF | + |
| TCTAAGATCACTG | 346570 | 346582 | *ORF240A* | ORF | + |
| TTGGAGAGTACAC | 346604 | 346616 | *ORF240A* | ORF | + |
| CCGCAAAACGCCA | 321212 | 321224 | *ORF240B* | ORF | + |
| TGTTAGATTGCAA | 320834 | 320846 | *ORF240B* | ORF | + |
| CCTTAAAGCAGTA | 270493 | 270505 | *ORF262* | ORF | - |
| CTCCAAAACGGAC | 359404 | 359416 | *ORF275* | ORF | + |
| CGGTAAATCAAGT | 359466 | 359478 | *ORF275* | ORF | + |
| TTCTAAAATGGTT | 84375 | 84387 | *ORF291* | ORF | - |
| CTGGAGAACGAGA | 69013 | 69025 | *ORF294* | ORF | - |
| TGCAAAATCAATA | 31159 | 31171 | *ORF315* | ORF | + |

| | | | | | |
|---|---|---|---|---|---|
| TCTAAAATCATAG | 222753 | 222765 | *rpl2 5'UTR* | 5'UTR | - |
| TCTAAGAGCGCCT | 222160 | 222172 | *rpl2 exon 1* | exon | - |
| TCTCAGATCGCTC | 222400 | 222412 | *rpl2 exon 1* | exon | - |
| CGTTAAAGTGACG | 220354 | 220366 | *rpl2 intron* | intron | - |
| TGGCAGAATGCTT | 220632 | 220644 | *rpl2 intron* | intron | - |
| TGGAAGATCATTT | 184883 | 184895 | *rpl5* | exon | - |
| TGGAAAATTGGCT | 185164 | 185176 | *rpl5* | exon | - |
| CTGAAGAACAAAA | 23125 | 23137 | *rps3 exon 2* | exon | + |
| TTCCAAAATAAGT | 22378 | 22390 | *rps3 exon 2* | exon | + |
| TTCTAAATCACTT | 22254 | 22266 | *rps3 exon 2* | exon | + |
| TGTCAAAGTGTGG | 23349 | 23361 | *rps3 exon 2* | exon | + |
| TGCTAAAATAGTG | 23160 | 23172 | *rps3 exon 2* | exon | + |
| TCGGAGAGCACTT | 21725 | 21737 | *rps3 intron* | intron | + |
| TCTCAAAGTATGG | 21622 | 21634 | *rps3 intron* | intron | + |
| CTGGAAAACACCC | 20366 | 20378 | *rps3 intron* | intron | + |
| CCGAAGATTGAGG | 160105 | 160117 | *rps4* | exon | + |
| CGCGAGAGTGTAG | 79360 | 79372 | *rps7 5'UTR* | 5'UTR | + |
| TTGGAGATTGAGG | 79213 | 79225 | *rps7 5'UTR* | 5'UTR | + |
| TGCGAAAGCATGG | 127454 | 127466 | *rrn18* | rRNA | - |
| CGGTAAAACGGGG | 127686 | 127698 | *rrn18* | rRNA | - |
| TGCCAAAACGGCT | 126757 | 126769 | *rrn18* | rRNA | - |
| TTTTAGAACAGAA | 140863 | 140875 | *rrn26* | rRNA | - |
| TTCCAAAGTGTTC | 143671 | 143683 | *rrn26* | rRNA | - |
| CGGCAAAATGACC | 141947 | 141959 | *rrn26* | rRNA | - |
| TGGCAAAATACGG | 143064 | 143076 | *rrn26* | rRNA | - |
| CGTGAAAACACCC | 126085 | 126097 | *rrn5* | rRNA | - |
| CTGAAGATTACAG | 171591 | 171603 | *trnfM* | tRNA | - |
| TGGTAGAGCATAG | 189302 | 189314 | *trnG* | tRNA | + |
| TGGTAGAGCATTG | 19944 | 19956 | *trnK.1* | tRNA | + |
| TGGTAGAGCATTG | 346470 | 346482 | *trnk.2* | tRNA | + |
| TGGTAGAGCGGTC | 43559 | 43571 | *trnN* | tRNA | + |
| TGCTAAATCGACA | 41519 | 41531 | *trnS.2* | tRNA | + |
| TGCTAAATCGACA | 321900 | 321912 | *trnS.4* | tRNA | - |
| CGGTAGAACGTGG | 298918 | 298930 | *trnW* | tRNA | + |

**Table S2**. Quantification data of protein profiles.

**a.** Relative accumulation of native organellar complexes in wild type (Col-0) and *ppr9* plants.

| **Genetic line** | **holo-CI** | | **CIII$_2$** | **CIV** | **CV** | |
|---|---|---|---|---|---|---|
| | **CA2** | **Nad9** | | | **AtpA** | **AtpB** |
| **Col-0** | 1.00 | 1.00 | 1.00 | 1.00 | 1.00* | 1.00* |
| ***ppr9 (homozygote, HmZ)*** | ---**/*** | --- ** | 3.15 | 5.55 | 3.53* | 3.14* |
| **Comp. (*ppr9/PPR9*)** | 0.51 | 0.36 | 0.47 | 0.32 | 0.79 | 0.46 |

* - appearance of an additional protein band of lower mass (postulated as the $F_1$-ATPase particle)
** - below limit of detection.
*** - appearance of partially assembled CI intermediates.

**b.** Relative accumulation of mitochondrial proteins in wild type and *ppr9* plants.

| | **Col-0** | ***ppr9* (HmZ)** | **Comp. *ppr9/PPR9*** |
|---|---|---|---|
| **Complex I** | | | |
| CA2 | 1.00 | 0.47 | 0.89 |
| NAD9 | 1.00 | 0.06 | 0.34 |
| **Complex III** | | | |
| RISP | 1.00 | 4.15 | 0.95 |
| **Complex IV** | | | |
| COX2 | 1.00 | 1.47 | 0.73 |
| **Complex V (ATP synthase)** | | | |
| AtpA | 1.00 | 1.63 | 0.84 |
| **Other organellar proteins** | | | |
| VDAC1-5 | 1.00 | 2.34 | 1.03 |
| AOX1a* | 1.00 | 25.23 | 1.00 |

* - appearance of an additional protein band of lower mass

**Table S3.** List of oligonucleotides used for the screening of *ppr9* mutants and for generating the GFP construct in this study.

| **Gene / Construct** | **Forward oligo (5'→3')** | **Reverse oligo (5'→3')** |
|---|---|---|
| *ppr9* | TCGCGTCTTCAACACTTCCA | CACTTCCTCCTCCCGAATGG |
| *LB1.3 (T-DNA)* | TCGCGTCTTCAACACTTCCA | ATTTTGCCGATTTCGGAAC |
| *PPR9-GFP* *(PPR9 NcoI F+ PPR9 SpeI R)* | CTCACCATGGCCCGAAGATTTTA CCGGAAGCCCTT | AGTCACTAGTCGACGGCAGGAGATTACGAT |

**Table S4.** List of RT-qPCR oligonucleotides designed to specific different exon-exon (analysis of mRNA and processed transcripts) and intron-exon regions (analysis of pre-RNAs) in Arabidopsis mitochondria.

a. List of oligonucleotides used for the analysis of the mRNA profiles in wild type and mutant plants

| gene target | oligo's name | sequence (5'→3') |
|---|---|---|
| *atp1* | *atp1F* | TCACTTCGACACGTCTTTGC |
| | *atp1R* | GGAATGGCCTTGAATCTTGA |
| *atp6* | *atp6-1F* | TCTTTTGCGAGTCAATGCAC |
| | *atp6-1R* | TCTCGCGTATCTCACATTGC |
| *atp8* | *atp8F* | CCGTCGACTTATTGGGAAAA |
| | *atp8R* | TTCCTTGGCCATGTACAACA |
| *atp9* | *atp9F* | CATTCCCTCTGACGTCGAAT |
| | *atp9R* | TCGTCGATTCTTACCCTCGT |
| *atp4* | *atp4F* | GGATCAGCTTGCGAATTTGT |
| | *atp4R* | GCAAATTGCTTCCCCACTAA |
| *ccmb* | *ccmBF* | TCTTGGAATCACATCCAGCA |
| | *ccmBR* | CGAGACCGAAATTGGAAAAA |
| *ccmc* | *ccmCF* | AGCTACGCGCAAATTCTCAT |
| | *ccmCR* | GCCGTGGCGATATAAACAAT |
| *ccmfc* | *ccmFcF* | CACATGGAGGAGTGTGCATC |
| | *ccmFcR* | GTGGGTCCATGTAAATGATCG |
| *ccmfn-1* | *ccmFN1F* | AGCTCTTGGCATTGCTTTGT |
| | *ccmFN1R* | AGTGCCACAATCCCATTCAT |
| *ccmfn-2* | *ccmFN2F* | CGTGTCGTTCGTAATGGAAA |
| | *ccmFN2R* | TGATAAGCCCACCAACTTCC |
| *cob* | *cobF* | TGCCGGAATGGTATTTCCTA |
| | *cobR* | GCCAAAAGCAACCAAAACAT |
| *cox1* | *cox1F* | GTAGCTGCGGTGAAGTAGGC |
| | *cox1R* | CTGCCTGGATTCGGTATCAT |
| *cox2* | *cox2F* | TGATGCTGTACCTGGTCGTT |
| | *cox2R* | TGGGGGATTAATTGATTGGA |
| *cox3* | *cox3F* | CCGTAACTTGGGCTCATCAT |
| | *cox3R* | AAACCATGAAAGCCTGTTGC |
| *mttb* | *mttBF* | GGGGTCTTTCTTTGGAAACC |
| | *mttBR* | TCTCCCTCATTCCACTCGTC |
| *nad1 exons 1-2* | *nad1 1-2F* | GACCAATAGATACTTCATAAGAGACCA |
| | *nad1 1-2R* | TTGCCATATCTTCGCTAGGTG |
| *nad1 exons 2-3* | *nad1 2-3F* | ATTCAGCTTCCGCTTCTGG |
| | *nad1 2-3R* | TCTGCAGCTCAAATGGTCTC |
| *nad1 exons 3-4* | *nad1 3-4F* | AAAAGAGCAGACCCCATTGA |
| | *nad1 3-4R* | TCCGTTTGATCTCCCAGAAG |
| *nad1 exons 4-5* | *nad1 4-5F* | AGCCCGGGATCTTCTTGA |
| | *nad1 4-5R* | TCTTCAATGGGGTCTGCTC |
| *nad2 exons 1-2* | *nad2 exons 1-2F* | GCGAGCAGAAGCAAGGTTAT |
| | *nad2 exons 1-2R* | GGATCCTCCCACACATGTTC |
| *nad2 exons 2-3* | *nad2 exons 2-3F* | AAAGGAACTGCAGTGATCTTGA |
| | *nad2 exons 2-3R* | AATATTTGATCTTAGGTGCATTTTC |
| *nad2 exons 3-4* | *nad2exons 3-4F* | GCGCAATAGAAAGGAATGCT |
| | *nad2exons 3-4R* | CTATGGGTCTACTGGAGCTACCC |
| *nad2 exons 4-5* | *nad2exons 4-5F* | CAAAGGAGAGGGGTATAGCAA |
| | *nad2exons 4-5R* | TATTTGTTCTTCGCCGCTTT |
| *nad3* | *nad3F* | CGAATGTGGTTTCGATCCTT |
| | *nad3R* | GCACCCCTTTTCCATTCATA |
| *nad4 exons 1-2* | *nad4 exons 1-2F* | ATTCTATGTTTTTCCCGAAAGC |

| | | |
|---|---|---|
| | *nad4 exons 1-2R* | GAAAAACTGATATGCTGCCTTG |
| *nad4 exons 2-3* | *nad4 exons 2-3F* | AATACCCATGTTTCCCGAAG |
| | *nad4 exons 2-3R* | TGCTACCTCCAATTCCCTGT |
| *nad4exons 3-4* | *nad4exons 3-4F* | TTCCTCCATAAATTCTCCGATT |
| | *nad4exons 3-4R* | TGAAATTTGCCATGTTGCAC |
| *nad4L* | *nad4L-F* | GGGGAATCCTCCTTAATAGACG |
| | *nad4L-R* | AACGAAAATGGCTAACCCAATA |
| *nad5 exons 1-2* | *nad5 exons 1-2F* | TGGACCAAGCTACTTATGGATG |
| | *nad5 exons 1-2R* | CCATGGATCTCATCGGAAAT |
| *nad5 exons 2-3* | *nad5 exons 2-3F* | TACCTAAACCAATCATCATATC |
| | *nad5 exons 2-3R* | CTGGCTCTCGGGAGTCTCTT |
| *nad5 exons 3-4* | *nad5exons 3-4F* | AACTCGGATTCGGCAAGAA |
| | *nad5exons 3-4R* | GATATGATGATTGGTTTAGGTA |
| *nad5 exons 4-5* | *nad5exons 4-5F* | AACATTGCAAAGGCATAATGA |
| | *nad5exons 4-5R* | GTTCCTGCGTTTCGGATATG |
| *nad6* | *nad6F* | TATGCCGGAAAGGTACGAAG |
| | *nad6R* | GTGAGTGGGTCAGTCGTCCT |
| *nad7 exons 1-2* | *nad7 exons 1-2F* | ACCTCAACATCCTGCTGCTC |
| | *nad7 exons 1-2R* | AAGGTAAAGCTTGAAGATAAGTTTTGT |
| *nad7 exons 2-3* | *nad7 exons 2-3F* | GAGGGACTGAGAAATTAATAGAGTACA |
| | *nad7 exons 2-3R* | TGGTACCTCGCAATTCAAAA |
| *nad7 exons 3-4* | *nad7exons 3-4F* | ACTGTCACTGCACAGCAAGC |
| | *nad7exons 3-4R* | CATTGCACAATGATCCGAAG |
| *nad7 exons 4-5* | *nad7exons 4-5F* | GATCAAAGCCGATGATCGTAA |
| | *nad7exons 4-5R* | AGGTGCTTCAACTGCGGTAT |
| *nad9* | *nad9F* | GGATGACCCTCGAAACCATA |
| | *nad9R* | CACGCATTCGTGTACAAACC |
| *rpl2* | *rpl2F* | CCGAAGACGGATCAAGGTAA |
| | *rpl2R* | CGCAATTCATCACCATTTTG |
| *rpl5* | *rpl5F* | AAGGGGTTCGACAGGAAAGT |
| | *rpl5R* | CGTATTTCGACCGGAAAATC |
| *rpl16* | *rpl16F* | GAGCATTTGCCAAACTCACA |
| | *rpl16R* | CGGACACTTTCATCGTGCTA |
| *rps3* | *rps3F* | CCGATTTCGGTAAGACTTGG |
| | *rps3R* | AGCCGAAGGTGAGTCTCGTA |
| *rps4* | *rps4F* | ACCCATCACAGAGATGCACA |
| | *rps4R* | TCACACAAACCCTTCGATGA |
| *rps7* | *rps7F* | CTCGAACTGAACGCGATGTA |
| | *rps7R* | AAGCTGCTTCAAGGATCCAA |
| *rps12* | *rps12F* | AGCCAAAGTACGGTTGAGCA |
| | *rps12R* | TTTGGGTTTTTCTGCACCAT |
| *matR* | *matR-F* | AATTTTTGCGAGAGCTGGAA |
| | *matR-R* | TTGAACCCCGTCCTGTAGAC |
| *rrn18* | *rrn18F* | CGTCACCTGGGTCAAAAACT |
| | *rrn18R* | GCTTGAAAACCGAAGTGAGC |
| *rrn26* | *rrn26F* | GACGAGACTTTCGCCTTTTG |
| | *rrn26R* | CTTGGAGCGAATTGGATGAT |
| *rrn5* | *rrn5F* | CCGACCTCGATATGTGGAATCGTC |
| | *rrn5R* | TGGACCATGTCTCCCGAACAATC |
| *18S rRNA* (At3g41768) | *18S nucl-F* | AAACGGCTACCACATCCAAG |
| | *18S nucl-R* | ACTCGAAAGAGCCCGGTATT |
| *ACTIN2* (At3g18780) | *actin2-F* | GGTAACATTGTGCTCAGTGGTGG |
| | *actin2-R* | AACGACCTTAATCTTCATGCTGC |
| *GAPDH* (At3g04120) | *GAPDH-F* | TCTCGATCTCAATTTCGCAAAA |
| | *GAPDH-R* | CGAAACCGTTGATTCCGATTC |

b. List of oligonucleotides designed for the analysis of the 23 organellar group II introns in Arabidopsis.

| Gene | Forward oligo (5'→3') | Reverse oligo (5'→3') |
|---|---|---|
| *rpl2* intron 1 exon 2 | TTAGGAAGAGCCGTACGAGG | CGCAATTCATCACCATTTTG |
| *rps3* intron 1 exon 2 | AGCCGAAGGTGAGTCTCGTA | TCTACGGCGGGGTCACTAT |
| *cox2* intron 1 exon 2 | TGGGGGATTAATTGATTGGA | AGCAGTACGAGCTGAAAGGC |
| *ccmFc* intron 1 exon 1 | CCCGGATCGAATCAGAGTT | CACATGGAGGAGTGTGCATC |
| *nad1* intron 1 exon 2 | GACCAATAGATACTTCATAAGAGACCA | CGTGCTCGTACGGTTCATAG |
| *nad1* intron 2 exon 2 | GGTTGGGTTAGGGGAACATC | TCTGCAGCTCAAATGGTCTC |
| *nad1* intron 3 exon 4 | AAAAGAGCAGACCCCATTGA | GGGAGCTGTATGAGCGGTAA |
| *nad1* intron 4 exon 5 | AGCCCGGGATCTTCTTGA | ACGGAGCTGCATCCCTACT |
| *nad2* intron 1 exon 2 | GCGAGCAGAAGCAAGGTTAT | CCCATTCCTAACCAGTGGAG |
| *nad2* intron 2 exon 2 | CCCGATCCGATAGTTTACAA | AATATTTGATCTTAGGTGCATTTTC |
| *nad2* intron 3 exon 4 | GCGCAATAGAAAGGAATGCT | GGCGAATTTCAAACTTGTGG |
| *nad2* intron 4exon 4 | CTTATTCGTGGCAACCTTCC | TATTTGTTCTTCGCCGCTTT |
| *nad4* intron 1 exon 2 | CCGTATGATGCGGAAGTCTC | GAAAAACTGATATGCTGCCTTG |
| *nad4* intron 2 exon 3 | GCGGAACGACCAGAAAAATA | TGCTACCTCCAATTCCCTGT |
| *nad4* intron 3 exon 4 | TCTAGCTTGGTTCGGAGAGC | TGAAATTTGCCATGTTGCAC |
| *nad5* intron 1 exon 2 | TGGACCAAGCTACTTATGGATG | TTCGCAAATAGGTCCGACT |
| nad5 intron 2-exon 2 | GTACGATCGTGTCGGGTGA | CTGGCTCTCGGGAGTCTCTT |
| nad5 intron 3-exon 4 | AACTCGGATTCGGCAAGAA | GCCGTGTAATAGGCGACCA |
| *nad5* intron 4 exon 5 | AACATTGCAAAGGCATAATGA | CCTGTAAACCCCCATGATGT |
| *nad7* intron 1 exon 2 | ACGGTTTTTAGGGGGATCTG | AAGGTAAAGCTTGAAGATAAGTTTTGT |
| *nad7* intron 2 exon 3 | AGTGGGAGAGCCGTGTTATG | TGGTACCTCGCAATTCAAAA |
| *nad7* intron 3 exon 4 | TAAAGTGAAGTGGTGGGCCT | CATTGCACAATGATCCGAAG |
| *nad7* intron 4 exon 5 | CGGCCAAATGACTACAGGAT | AGGTGCTTCAACTGCGGTAT |

**Table S5.** List of antibodies used for the analysis of organellar proteins in wild type and *ppr9* mutant plants.

| Antibody | Protein I.D. | origin | serum | dilution | Reference / source |
|---|---|---|---|---|---|
| AOX1a | Ubiquinol (Alternative) oxidase 1a | *Arabidopsis thaliana* | Rabbit (polyclonal) | 1/1,000 | Agrisera antibodies, AS04 054 |
| AtpA | Mitochondrial ATP-synthase subunit 1 (A) | *Nicotiana tabacum* | Mouse (monoclonal) | 1/500 | Thomas Elton collection |
| AtpB | Mitochondrial ATP-synthase subunit 2 (B) | *Nicotiana tabacum* | Mouse (monoclonal) | 1/500 | Thomas Elton collection |
| CA2 | g-carbonic anhydrase-like subunit 2 | *Arabidopsis thaliana* | Rabbit (polyclonal) | 1/1,000 | (Perales *et al.* 2005),(Sunderhaus *et al.* 2006) |
| COX2 | Cytochrome C oxidase subunit-2 | *Nicotiana tabacum* | Mouse (monoclonal) | 1/100 | Thomas Elton collection |
| NAD9 | NADH-dehydrogenase complex subunit-9 | *Triticum spp.* | Rabbit (polyclonal) | 1/50,000 | (Lamattina et al., 1993) |
| RISP | Rieske iron-sulfur protein | *Arabidopsis thaliana* | Rabbit (polyclonal) | 1/5,000 | Gift of Prof. Ian Small, UWA |
| Porin/VDAC | Voltage-dependent anion-selective channel protein 1-5 | *Arabidopsis thaliana* | Rabbit (polyclonal) | 1/1,000 | Agrisera antibodies, AS07 212 |

**Perales, M., Eubel, H., Heinemeyer, J., Colaneri, A., Zabaleta, E. and Braun, H.-P.** (2005) Disruption of a nuclear gene encoding a mitochondrial gamma carbonic anhydrase reduces complex I and supercomplex I+III2 levels and alters mitochondrial physiology in Arabidopsis. *J Mol Biol*, **350**, 263-277.

**Sunderhaus, S., Dudkina, N.V., Jansch, L., Klodmann, J., Heinemeyer, J., Perales, M., Zabaleta, E., Boekema, E.J. and Braun, H.P.** (2006) Carbonic anhydrase subunits form a matrix-exposed domain attached to the membrane arm of mitochondrial complex I in plants. *J Biol Chem*, **281**, 6482-6488.

**a**

predicted cleavage site

MRRFYRKPFSVPLLNRPHCSSQSHCLYKNGDFLSDDSKCSPLSSSRTSVRWVF | NSSSLPPPEWIEPFNDVSDLVKSNRNLLPSPWV
SQILNLLDGSASMESNLDGFCRKFLIKLSPNFVSFVLKSDEIREKPDIAWSFFCWSRKQKKYTHN | **LECYVSLVDVLALAKDVDRIR**
**FVSSEIKKFEFPMT | VSAANALIKSFGKLGMVEELLWVWRKMKENGIEPT | LYTYNFLMNGLVSAMFVDSAERVFEVMESGRIKPD |**
**IVTYNTMIKGYCKAGQTQKAMEKLRDMETRGHEAD | KITYMTMIQACYADSDFGSCVALYQEMDEKGIQVP | PHAFSLVIGGLCKEG**
**KLNEGYTVFENMIRKGSKPN | VAIYTVLIDGYAKSGSVEDAIRLLHRMIDEGFKPD | VVTYSVVVNGLCKNGRVEEALDYFHTCRFD**
**GLAIN | SMFYSSLIDGLGKAGRVDEAERLFEEMSEKGCTRD | SYCYNALIDAFTKHRKVDEAIALFKRMEEEEGCDQT | VYTYTILL**
**SGMFKEHRNEEALKLWDMMIDKGITPT | AACFRALSTGLCLSGKVARACKILDELAPMGVILD | AACEDMINTLCKAGRIKEACKLA**
**DGITERGREVP | GRIRTVMINALRKVGKADLAMKLMHSKIGIGYERM |** GSVKRRVKFTTLLETCFDS

**b**

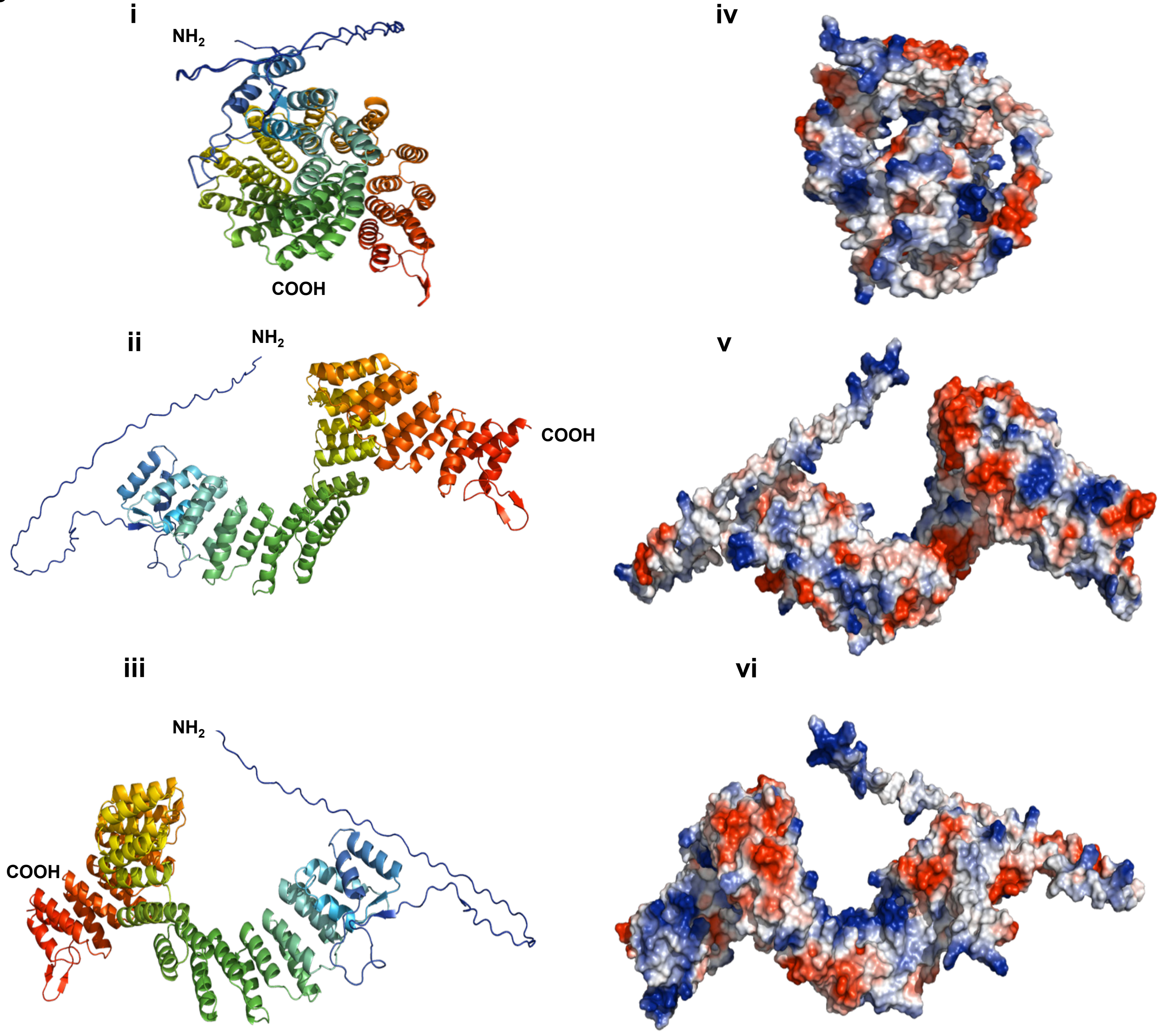


Figure S1

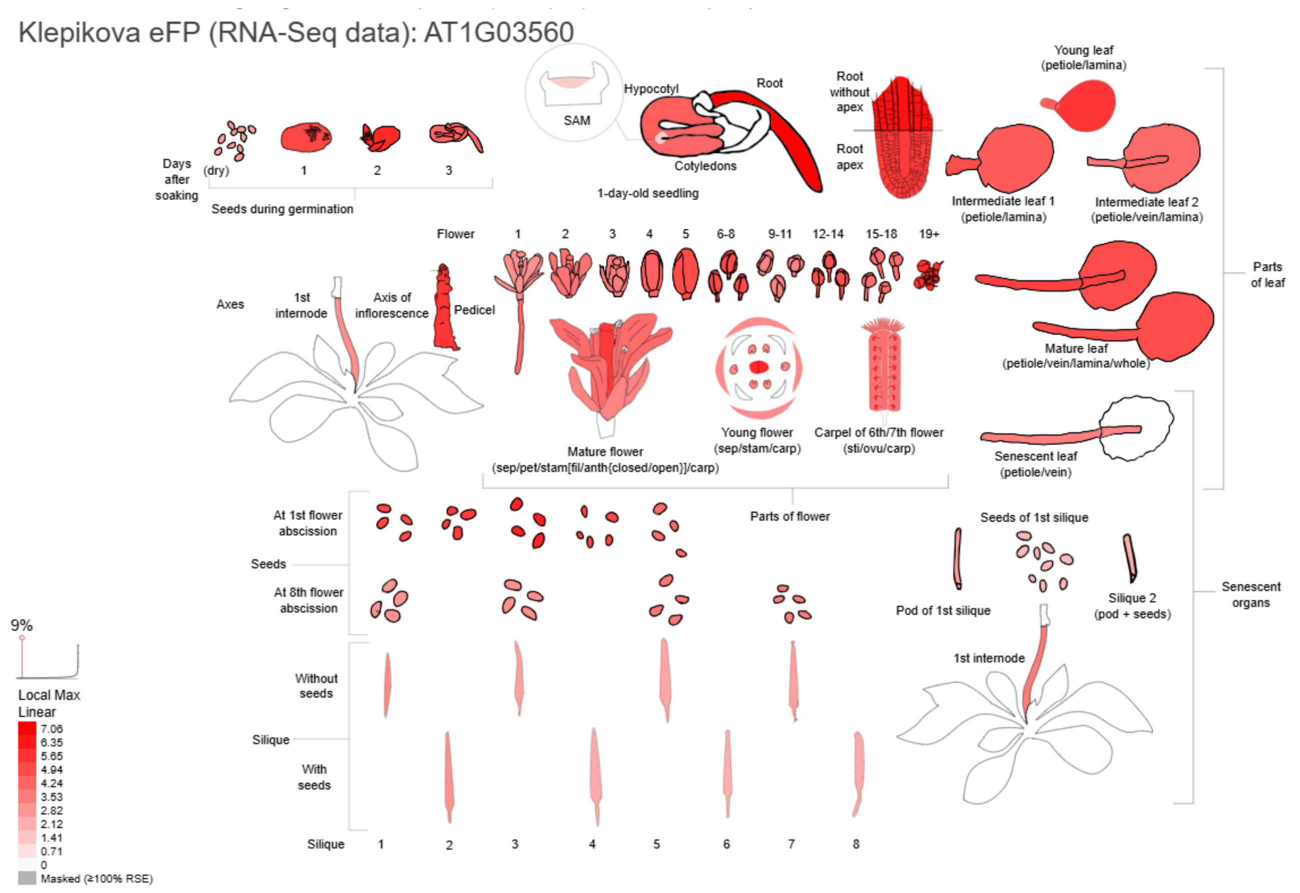

Klepikova eFP (RNA-Seq data): AT1G03560
SAM
Hypocotyl
Root
Root without apex
Root apex
Cotyledons
1-day-old seedling
Young leaf (petiole/lamina)
Intermediate leaf 1 (petiole/lamina)
Intermediate leaf 2 (petiole/vein/lamina)
Days after soaking
(dry)
1
2
3
Seeds during germination
Flower
1
2
3
4
5
6-8
9-11
12-14
15-18
19+
Axes
1st internode
Axis of inflorescence
Pedicel
Parts of leaf
Mature leaf (petiole/vein/lamina/whole)
Mature flower (sep/pet/stam[fil/anth{closed/open}]/carp)
Young flower (sep/stam/carp)
Carpel of 6th/7th flower (sti/ovu/carp)
Senescent leaf (petiole/vein)
At 1st flower abscission
Seeds
At 8th flower abscission
Parts of flower
Seeds of 1st silique
Pod of 1st silique
Silique 2 (pod + seeds)
Senescent organs
1st internode
9%
Local Max
Linear
7.06
6.35
5.65
4.94
4.24
3.53
2.82
2.12
1.41
0.71
0
Masked (≥100% RSE)
Without seeds
Silique
With seeds
Silique
1
2
3
4
5
6
7
8


Figure S2

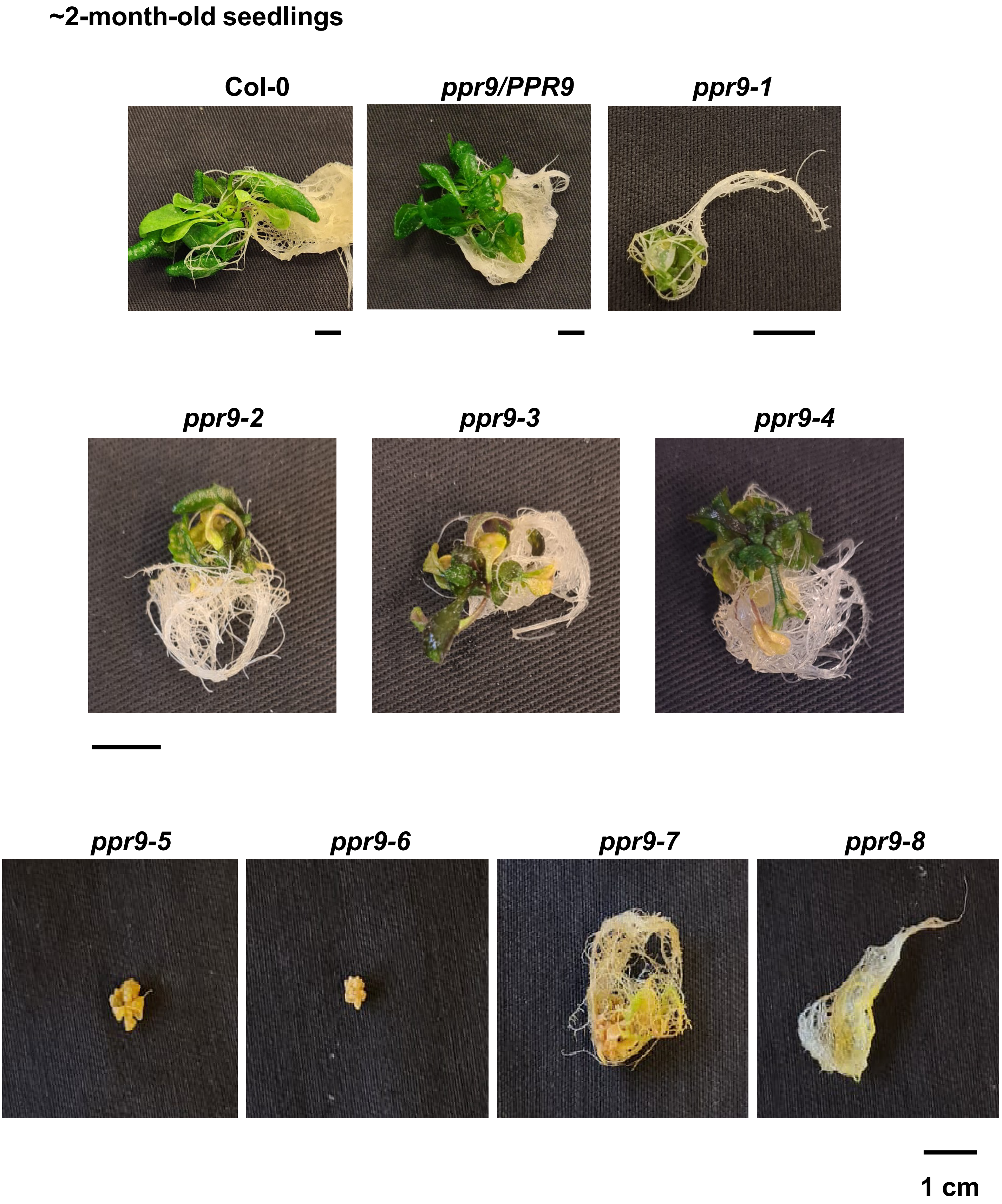
~2-month-old seedlings
Col-0
ppr9/PPR9
ppr9-1
ppr9-2
ppr9-3
ppr9-4
ppr9-5
ppr9-6
ppr9-7
ppr9-8
1 cm

**Figure S3**

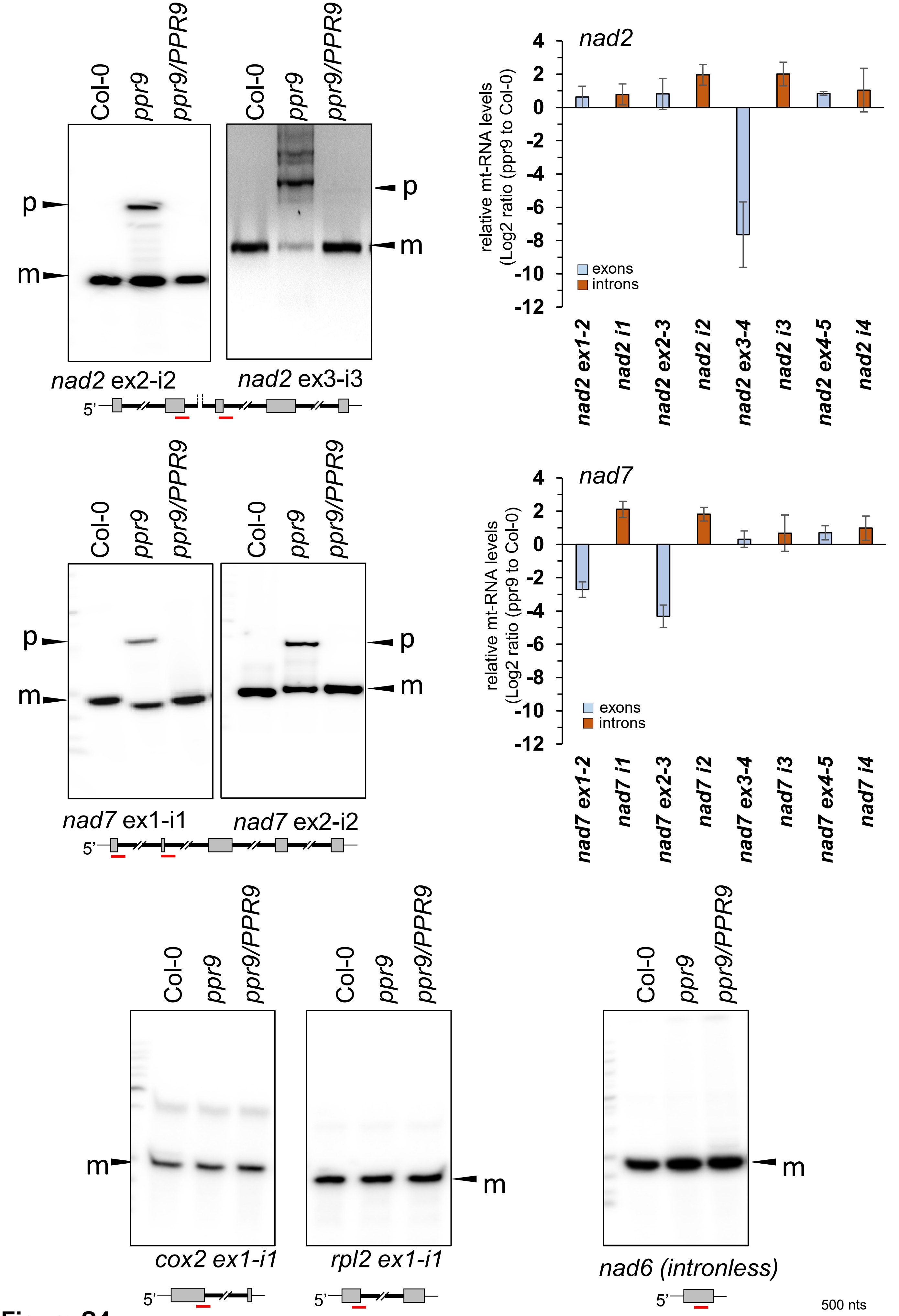

Col-0
ppr9
ppr9/PPR9
p
m
nad2 ex2-i2
nad2 ex3-i3
5'
nad2
relative mt-RNA levels
(Log2 ratio (ppr9 to Col-0)
4
2
0
-2
-4
-6
-8
-10
-12
exons
introns
nad2 ex1-2
nad2 i1
nad2 ex2-3
nad2 i2
nad2 ex3-4
nad2 i3
nad2 ex4-5
nad2 i4
nad7 ex1-i1
nad7 ex2-i2
nad7
nad7 ex1-2
nad7 i1
nad7 ex2-3
nad7 i2
nad7 ex3-4
nad7 i3
nad7 ex4-5
nad7 i4
cox2 ex1-i1
rpl2 ex1-i1
nad6 (intronless)
500 nts


**Figure S4**
